# Modeling Adaptive and Non-adaptive Responses of Populations to Environmental Change

**DOI:** 10.1101/090894

**Authors:** Tim Coulson, Bruce E Kendall, Julia Barthold, Floriane Plard, Susanne Schindler, Arpat Ozgul, Jean-Michel Gaillard

## Abstract

Understanding how the natural world will be impacted by environmental change over the coming decades is one of the most pressing challenges facing humanity. Addressing this challenge is difficult because environmental change can generate both population level plastic and evolutionary responses, with plastic responses being either adaptive or non-adaptive. We develop an approach that links quantitative genetic theory with data-driven structured models to allow prediction of population responses to environmental change via plasticity and adaptive evolution. After introducing general new theory, we construct a number of example models to demonstrate that evolutionary responses to environmental change over the short-term will be considerably slower than plastic responses, and that the rate of adaptive evolution to a new environment depends upon whether plastic responses are adaptive or non-adaptive. Parameterization of the models we develop requires information on genetic and phenotypic variation and demography that will not always be available, meaning that simpler models will often be required to predict responses to environmental change. We consequently develop a method to examine whether the full machinery of the evolutionarily explicit models we develop will be needed to predict responses to environmental change, or whether simpler non-evolutionary models that are now widely constructed may be sufficient.

## Introduction

Ecosystems from the deep ocean to the high arctic, from deserts to tropical forests are responding to environmental change. Understanding and predicting these responses is one of the most pressing issues currently facing humanity. For this reason, in the last quarter of a century, there has been considerable interest in developing ways to understand how the natural world will be affected by environmental change (Bossdorf et al., 2008; Dawson et al., 2011; Gilbert and Epel, 2009; Hoffmann and Sgrò, 2011; Ives, 1995; Lavergne et al., 2010; Wiens et al., 2009). We introduce a new, general approach combining insights from structured population modeling and evolutionary genetics that allows us to examine how adaptive evolution and plasticity contribute to the way that populations, and consequently the ecosystems in which they are embedded, respond to environmental change.

In order to understand how evolution and plasticity contribute to population responses to environment change it is necessary to appreciate how different levels of biological organization – alleles, genotypes, phenotypes, populations – are linked, as well as feedbacks between the different levels. First, evolution is defined as a change in allele frequencies (Charlesworth, 1994). Allele frequencies change as a direct consequence of changes in the frequencies of the genotypes the alleles occur in, and genotype frequencies can change with a change in the distribution of the phenotypes they code for (Fisher, 1930). The dynamics of phenotypic trait distributions are determined by differential birth, death, development and inheritance rates across phenotypic trait values, where inheritance is defined in the broad sense as the map between parental and offspring phenotypes (Easterling et al., 2000; Rees et al., 2014). Given these links between different levels of biological organization, there can be a cascading dynamic at the level of the phenotype, the genotype and the allele caused by differences in the demography of individuals with different phenotypic trait values (Lynch and Walsh, 1998). Another consequence of this variation is the ecology of the system: population dynamics are an emergent property of who lives, who breeds and with whom, as are the dynamics of the community and ecosystem the population is embedded within (?).

Although the cascading ecological and evolutionary consequences of variation in demographic rates is relatively straightforward to grasp, the devil is in the detail, and in particular how alleles combine to make genotypes, how genotypes influence phenotypes, and how phenotypes influence demographic rates (Coulson et al., 2011). The rate and direction of evolution depends upon how these links influence the relative fitness of each allele within the population (Charlesworth, 1994). The challenge is these links are often complicated, particularly for complex phenotypic traits like body size that are routinely measured by field biologists. The complexity arises not only because large gene networks and multiple cell types can contribute to the phenotype, but also because the environment makes a contribution too via plasticity defined as change in a phenotypic trait distribution that is not caused by genetic change (Gavrilets and Scheiner, 1993; Lande, 2009; ?).

The environment can be partitioned into biotic and abiotic components (?). The biotic component captures the sizes and structures of the population of the focal species and of all other species with which it interacts. The abiotic environment includes weather, mineral and water available. The biotic and abiotic environments can influence one another, although the influence of the biotic environment on the abiotic environment typically plays out over geological time scales (one exception being manmade climate change).

The biotic and abiotic environment can influence both the map between genotype and phenotype (?), and between phenotype and demographic rates (?). Put another way, demographic rates are a function of phenotype-by-environment interactions, and phenotypic traits are a function of genotype-by-environment interactions. For quantitative phenotypic traits, genotype-by-environment interactions can often usefully be understood by treating the phenotype as consisting of a genetic and an environmental component, with the environmental component determined by aspects of the current and past biotic and abiotic environments (Falconer, 1960; Lande, 1982; ?). The environmental component of the phenotype can capture phenotypic change caused by individuals altering their physiology, metabolism, behavior or levels of gene expression. We use the term epigenetic to refer to any process that does not involve genetic change that is captured by the dynamics of the environmental component of the phenotype. The biotic and abiotic environment can also influence the generation of new alleles via, for example, retroviral insertions into the germline of their hosts, or via ultraviolet radiation (??). In Figure 1(a) we depict how different levels of biological organization are linked and feedback to influence one another.

**Figure 1.**
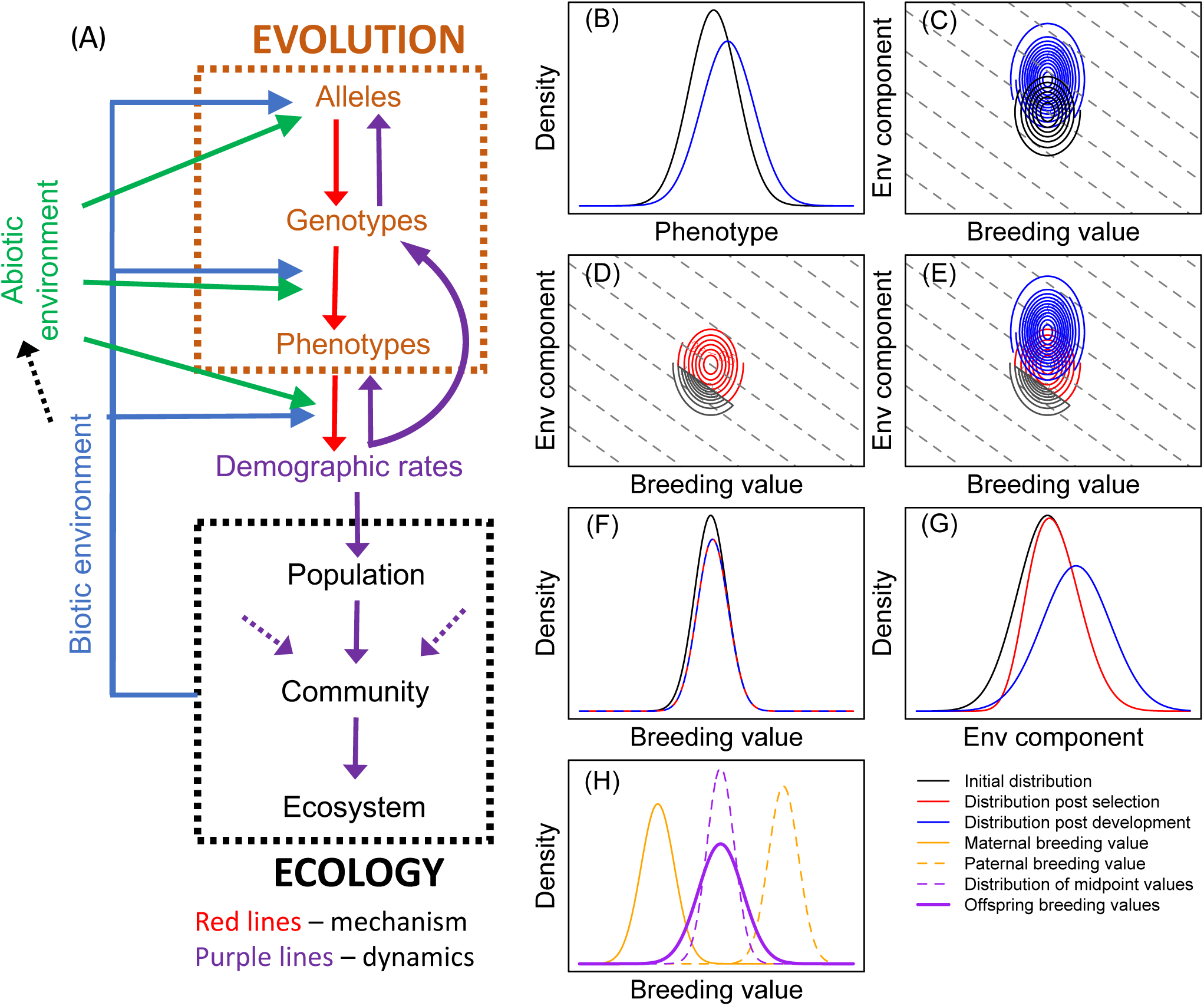
(A) linkages and feedbacks in biology. Evolution is defined as change in allele frequencies but is often inferred from the dynamics of genotypes and phenotypes. Research into links between alleles and genotypes, and particularly between genotypes and phenotypes often focuses on mechanism (red arrows). Differential survival and reproduction and patterns of mating determine (i) the dynamics of phenotypes, genotypes and alleles and (ii) population, community and ecosystem dynamics (purple arrows). Ecological dynamics determine the biotic environment that, along with the abiotic environment, can influence the generation of new alleles as well as the maps between genotype and phenotype and between phenotype and demographic rates. (B) IPMs track the dynamics of the phenotype distribution from t (black line) to t + 1 (blue line). (C) In our approach, we treat the phenotype as a bivariate distribution of an additive genetic (breeding value) and environmental component and iterate this distribution forwards. Dashed gray lines are clines, where each point on a cline denotes the same phenotypic trait value. There are two steps to iterate the phenotype forwards within a cohort. First (D) viability selection. In this example, all individuals with a trait value below a threshold have lower survival than those above the threshold, and second (E) development among survivors. Breeding values do not change within individuals as they age, meaning (F) only selection can generate change in the breeding value distribution within a cohort. In contrast, (G) selection and development can alter the distribution of the environmental component of the phenotype. The dynamics of the two components combine to generate the dynamics of the phenotype. (H) Mechanistic inheritance rules generate the distribution of offspring breeding value given parental breeding values in two steps. First, a distribution of mid-point parental breeding values is generated, before segregation variance is added to create a distribution of offspring breeding values. The inheritance rules for the map between the parental and environmental component of the phenotype are less constrained than genetic inheritance and are not shown.

How can this view of biology be used to inform how populations respond to environmental change? Environmental change occurs when the biotic or abiotic environment changes. Biotic changes can result from the arrival of a new species or an extinction within the ecosystem, or from evolution. In order to capture such change, and to model the links between alleles and demographic rates described above and in Figure 1(a), it is necessary for models to incorporate (i) the genotype-phenotype map at birth, (ii) how the phenotype develops, (iii) how the phenotype influences survival at each developmental stage, (iv) the population’s mating system and (v) patterns of mate choice based on the phenotype, as well as how these mate choice patterns influence (vi) reproductive success, (vii) the distribution of genotypes among offspring and (viii) how all these processes result in change in allele frequency and population size from one generation to the next. Processes (i) to (vi) (and consequently also (viii)) can be influenced by the biotic or abiotic environment. Integral Projection Models (IPMs) provide a very flexible structured modeling framework that allow each of these processes to be simultaneously modeled (Coulson, 2012; Easterling et al., 2000; Merow et al., 2014).

IPMs project the dynamics of phenotype distributions as a function of expected survival and reproduction, the way the phenotype develops and the distribution of offspring phenotypes (Easterling et al., 2000). Numerous quantities of interest to ecologists and evolutionary biologists describing life history, population dynamic and phenotypic traits can be calculated from IPMs (Childs et al., 2003a; Coulson et al., 2011, 2010; Ellner and Rees, 2006; Rees et al., 2014; Steiner et al., 2014, 2012; Vindenes and Langangen, 2015). They consequently offer great potential to study ecological and evolutionary responses to environmental change (Coulson et al., 2011). However, most IPMs to date have been restricted to phenotypic variation in that they do not include genotype-phenotype maps (Merow et al., 2014). A small number of evolutionarily explicit IPMs that do include these maps have been developed. For example, Coulson et al. (2011) used IPMs to track the distribution of body size and coat color in wolves, where coat color was determined by genotype at a single bi-allelic locus. Barfield et al. (2011) and Childs et al. (2016) developed IPMs of quantitative characters determined by a large number of unlinked loci of small effect. However, none of these models incorporates plasticity, nor different genetic influences on the phenotype at different ages, and these omissions limit their utility in predicting how populations will be influenced by environmental change (Chevin, 2015).

The aim of this paper is to introduce a general framework to allow prediction of how populations respond to environmental change. We do this by developing IPMs of the bivariate distribution of a phenotype split into its genetic and environmental components. The models incorporate different development and inheritance rules for each component of the phenotype. We develop and illustrate our framework using simple models. Our models reveal new insights into the way that plasticity can influence evolution, while also allowing us to retrieve key findings from evolutionary genetics that are already known.

## Methods and Results

### Modeling approach

We refer to functions as *f* (…) where the dots inside parentheses define the variables the function *f* operates on. Parameters of a function are referenced by the same letter as the function, with subscripts defining the variable they influence. For example, a parameter 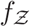 represents a parameter of function *f* that operates on variable 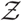. We reserve *I* for the intercept of functions and *a* for age. Age is only included in models for species with overlapping generations. Following standard convention for IPMs (Coulson, 2012; Merow et al., 2014; Rees et al., 2014; ?), we use primes to indicate the character value of an individual at the end of a time step. For example, this allows us to show how an individual with phenotype 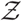 can develop over a time step to a potentially different phenotype 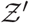, or how a parent with genotype 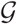 can produce an offspring with genotype 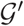. There are, of course, other notational conventions that could achieve the same objective, and we recognize that primes are used differently in evolutionary genetics; our notation is chosen to make clear how evolutionary processes can be included in the IPM framework. The definitions of all variables and functions are summarized in Table 1.

**Table 1:**
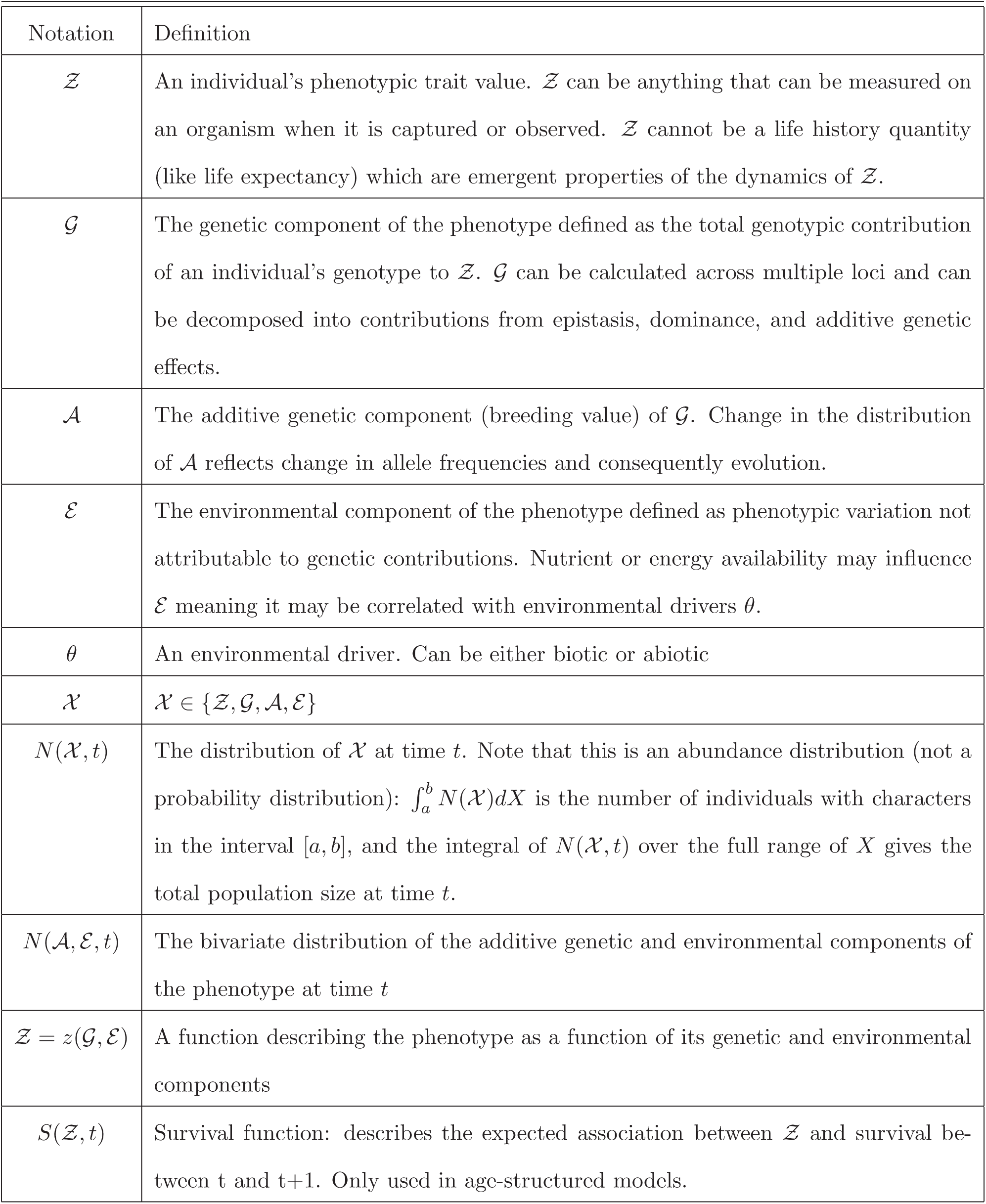
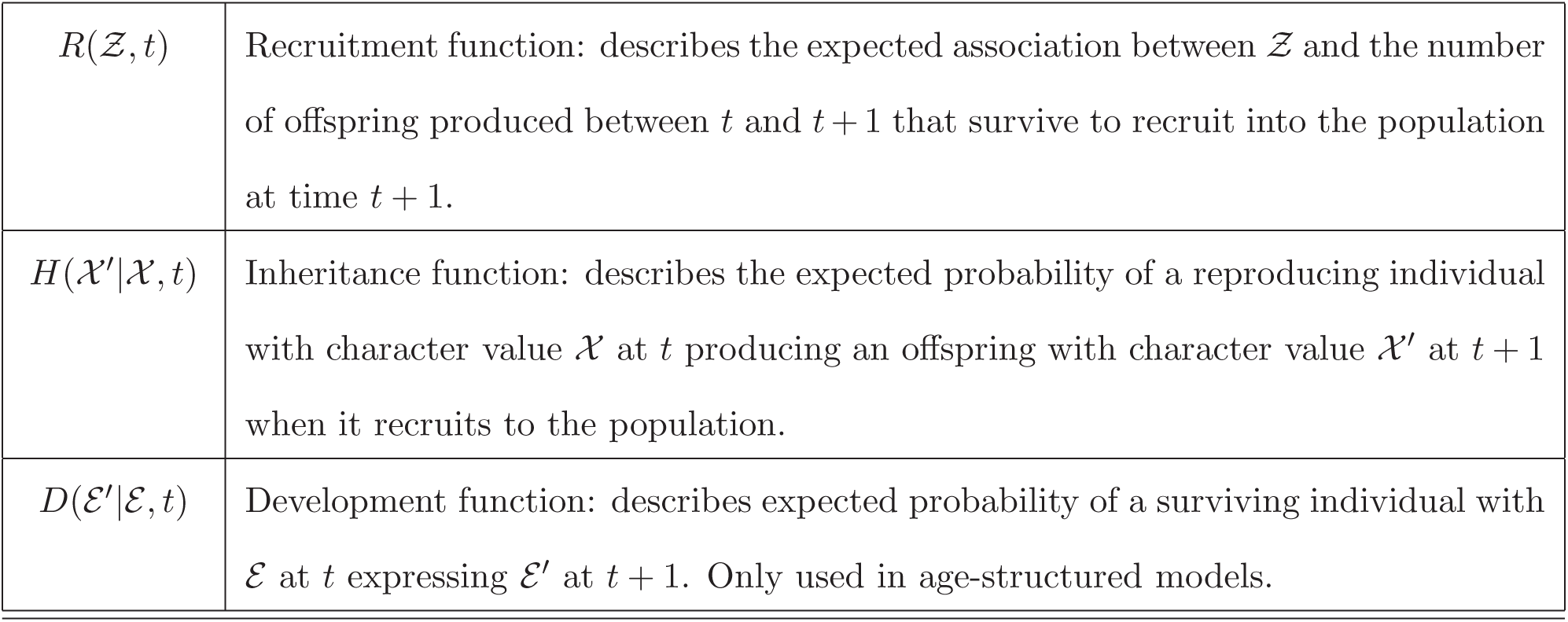
Notation used in the paper.

Our starting point is a widely used phenotypic modeling approach that many readers will be familiar with (Coulson, 2012; Merow et al., 2014; Rees et al., 2014). We then extend this approach by developing dynamic models of the phenotype decomposed into its genetic and environmental components. We start with a two-sex IPM that captures all demographic processes that can contribute to the dynamics of phenotypes – survival, recruitment, development, inheritance, and mating patterns (Coulson et al., 2011; Schindler et al., 2015; Traill et al., 2014a; ?) and which iterates forwards the distribution of the phenotype at time 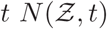 (Figure 1(B)).

The model consists of two equations – one for females and one for males – with each equation consisting of two additive components (?). The first component deals with survival and development of individuals already within the population, the second component deals with reproduction and the generation of phenotypes among newborns entering the population. We assume a pre-breeding census such that survival occurs before development and recruitment before inheritance, 
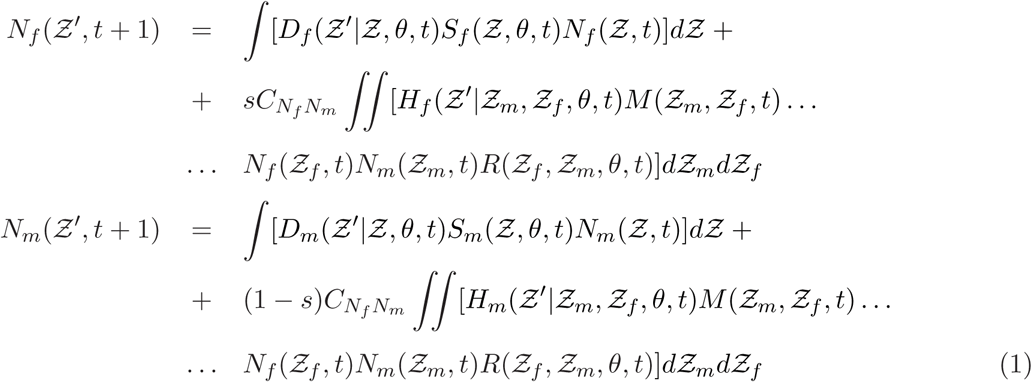
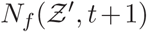 and 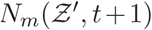 are distributions of phenotypes 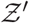 in respectively females and males at time *t* + 1; 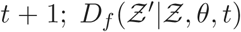 and 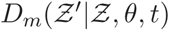 are the probability of the phenotype developing from 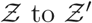 in respectively females and males between *t* and *t* + 1 as a function of environmental drivers *θ*;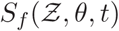 and 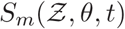 are survival functions for females and males from *t* to *t* + 1 including effects of phenotype and environmental drivers *θ*; *s* is the birth sex ratio measured as the proportion of female offspring produced; 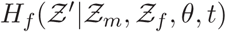 and 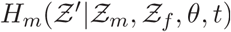 describe the probabilities of parents with phenotypes 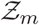 and 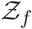 respectively producing male and female offspring with phenotype 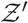 as a function of environmental drivers *θ* at time 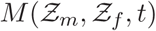 captures the rate of mating between a male with phenotype 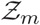 and a female with phenotype 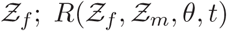 is the expected litter size given a mating between a male and a female with phenotypes 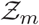 and 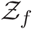 in environment *θ* at time *t*; *C_N_f_N_m__* is a normalization constant that is used to specify the mating system. In theory it could be combined with the mating function, but we follow the notation of ?. *C_N_f_N_m__* can be used to capture a range of mating systems. For example, if we follow ? and write, 
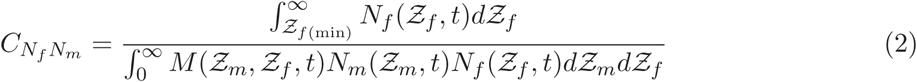
this adds a minimum size at which females can reproduce 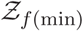. Depending on the mating behavior of the species, *C_N_f_N_m__* can be modified in various ways. For example, it can easily be altered such that the number of birth events is determined by the number of the rarer sex, as in monogamous species. Mate choice can be influenced by specifying different functions for 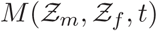. ? demonstrate how it can be specified for random mating, assortative mating, disassortative mating and size-selective mating.

In phenotypic IPMs, the phenotypic development functions are usually Gaussian probability functions (Easterling et al., 2000), e.g.: 
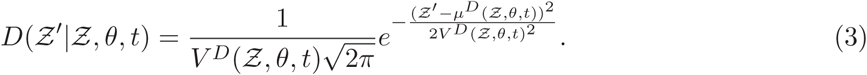

The functions 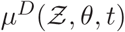 and 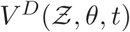 respectively describe the expected value of 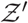 given 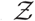 and *θ* at time *t* and the standard deviation around 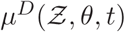. The Gaussian form can also be used for inheritance functions 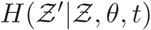 with functions *μ^Η^*(…) and *V^H^*(…).

The two-sex IPM described above is not evolutionarily explicit as it does not include mechanistic rules for genetic inheritance. We now take this phenotypic model and extend it to be evolutionarily explicit. We do this by writing the phenotype as a function of genetic 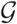 and environmental 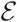 components 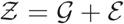. We assume that 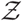 is a quantitative phenotype (i.e. measured in integer or real values). The genotypic value 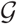 and environmental value 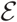 describe the numerical contributions of the genetic and environmental components of the phenotype to an individual’s phenotypic trait value. A simple map can consequently be written 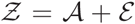 (Falconer, 1960).

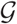 is determined by genotype, *g*. When the map between *g* and 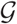 is additive, the dynamics of *g* and 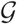 are identical (Falconer, 1960). This means that the dynamics of alleles are identical to the dynamics of genotypes in which they occur. In contrast, when alleles interact, either at a locus (dominance) or across loci (epistasis) the map between *g* and 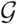 is not additive, and the dynamics of 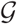 are not identical to the dynamics of *g* (Fisher, 1930). In classical quantitative genetics it is assumed that the map between *g* and 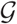 is additive (Falconer, 1960). Under these assumptions, it is not necessary to track the dynamics of *g* but evolution can be investigated by modeling the dynamics of just 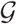. When the map is additive we refer to the genetic component of the phenotype as a breeding value and denote it 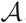.

In classical population genetics, when the contribution of dominance and epistasis to evolution are often a key focus, it is necessary to track the dynamics of *g* and calculate 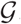 from each *g*. The map between 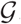 and the phenotype 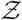 is often assumed to be one-to-one (?). In contrast, in quantitative genetics, the environment can influence the map between 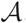 and 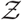 by influencing the value of the environmental component of the phenotype, 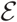 (Falconer, 1960). 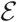 can take different values in different individuals and can vary within individuals throughout life. The dynamics of the phenotype may not consequently represent the dynamics of the genotypic value 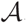. Statistical quantitative genetics is concerned with estimating moments of 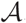 from 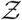 by correcting for environmental and individual variables that determine 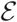 (Kruuk et al., 2008).

The genotype-phenotype map for phenotypic traits measured by biologists in free-living populations is rarely known, and quantitative genetic assumptions are widely adopted (Kruuk et al., 2008). In particular, the infinitesimal model is assumed in which 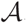 is determined by a large number of unlinked loci of small, additive, effect (Fisher, 1930). Until we have a better understanding of the genetic architecture of complex traits, this approach is the most powerful available to investigate evolution in the wild (Kruuk et al., 2008). We consequently adopt it here.

We track the joint distribution of the two components 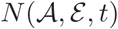. The utility of this is we can write expressions to describe the dynamics of each of the components separately, if necessary, before easily combining them to retrieve the dynamics of the phenotype. For 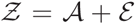 we can use a convolution (represented by the mathematical operator *) between the two components of the phenotype to construct the phenotype (Barfield et al., 2011).

Phenotypic plasticity and epigenetic inheritance are captured in the dynamics of 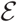. In previous quantitative genetic IPMs 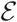 is a randomly distributed variable that captures developmental noise (Barfield et al., 2011; Childs et al., 2016). A key contribution of this paper is to show how 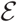 can be extended to also capture the biotic or abiotic environment as well as signatures of parental 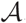 s and 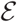 s. 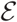 is defined as function of these drivers, and we write 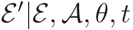 to capture the effects of 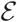, 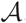 and the environment *θ* at time *t* on 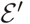.

We now expand terms in our two-sex phenotypic IPM to include the genotype-phenotype map 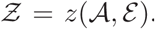. We start with the bivariate distribution of 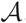 and 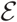 at time *t* among females that are already within the population at time *t*: 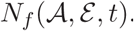. Viability selection now operates on this distribution. Viability selection is a simple multiplicative process describing the expected survival from *t* to *t* + 1 as a function of the phenotype. We can consequently write, 
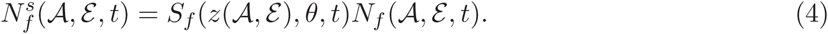

When it comes to development, 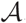 remains fixed throughout life while 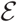 may vary, 
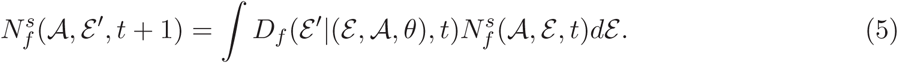

Recruitment is dealt with in a similar way to survival in that it is a multiplicative process, 
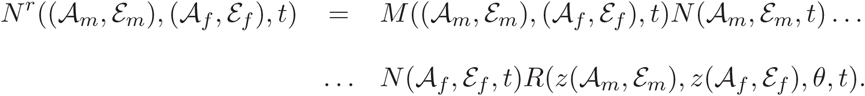

Note this is a recruitment related term of both male and female offspring that is not yet scaled by the normalization factor *C_N_f_N_m__*.

As with development, inheritance of the genetic and environmental components of the phenotype operates in different ways. For example, once mating pairs have formed and the number of offspring from each mating has been determined, the distribution of offspring genotypes is predictable. We can write the inheritance function for the genetic and environmental components of the phenotype as, 
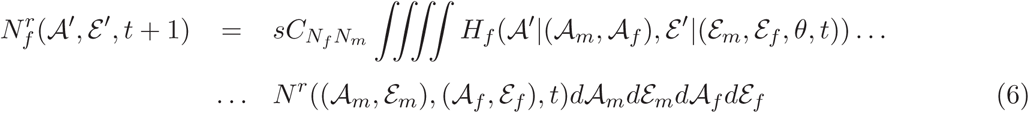
then, 
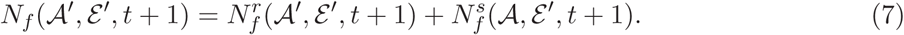

The same logic applies to the production of male offspring. We can construct the phenotype from the two components 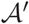 and 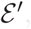, e.g. 
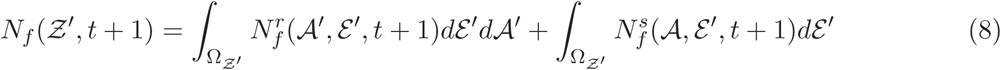
where 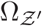 is the set of 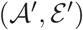 values satisfying 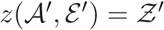. For the second integral in equation (8) we have 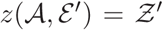 as the 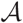 does not change within individuals and consequently has no prime.

The additivity assumption means that models of clonal inheritance can generate very similar predictions to models of two sexes, particularly if both males and females have similar demography. However, clonal models are simpler than two sex models (Lande, 1982). We utilize this consequence of the additivity assumption and initially work with clonal reproduction to examine how the dynamics of 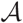 and 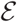 influence population and phenotypic trait dynamics and adaptive evolution. We can write a clonal model, 
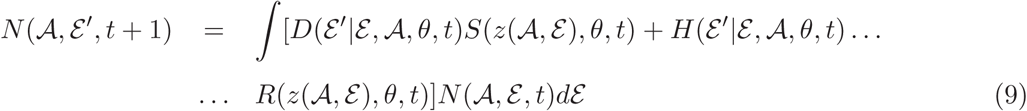
and 
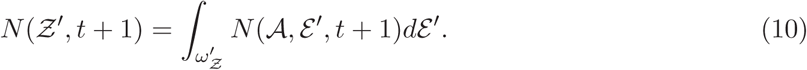

The above equations describe how the dynamics of a bivariate distribution of the genetic and environmental components of the phenotype. Figures 1(B-G) provide graphical examples of how these functions alter the bivariate distribution, and in particular how development and inheritance rules differ between the environmental and additive genetic components. To demonstrate these differences we now focus on developing univariate models of (i) 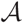, and (ii) 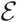. These models capture limits where all phenotypic variation among individuals is determined by (i) genetic variation and (ii) variation in the environmental component of the phenotype. We then combine insights from these univariate model and construct models of the bivariate distribution of 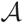 and 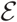.

We primarily work with linear functions for three reasons. First, they are easier to interpret and analyze than non-linear or non-additive forms. Second, when the environment changes impacting populations, responses, at least in the short term, can be well described with linear or linearized additive models (Cooch et al., 2001). Third, selection, the underpinning of evolution, is often directional and well described with linear or linearized associations between phenotypic traits and components of fitness (Kingsolver et al., 2001). Parameters used for all models are provided in Appendix A (§1.1), as are expressions to calculate key statistics used to show ecological and evolutionary change from model outputs (§1.2). Code to produce each figure is available on GitHub – https://github.com/tncoulson/QG-meets-IPM-figure-code/tree/master and Dryad (?).

### Adaptive Evolution

In this section we start with a simple clonal model of a univariate distribution of 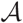. We go on to show how genetic constraints can be imposed to slow, or stop, evolution. We then extend this clonal model in two ways: first, to include a multivariate, age-structured, distribution of 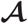, and second we relax the clonality assumption and compare the dynamics of clonal and sexual models. Finally, we introduce a new approximation to describe sexual reproduction and compare its performance with our initial approach.

Genotypes (and hence 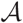) are determined at birth and remain fixed throughout life; neither are influenced by the environment. A consequence of this is the development function simplifies to a one-to-one map and can be removed from equation (5). We also start by considering clonal reproduction, which means that the inheritance function can also be removed as offspring genotype is identical to parental genotype. The dynamics of 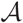 are consequently determined by the survival and reproduction functions – selection. In these models, as long as there is genetic variation within a population, and fitness is a monotonic function of genotype, evolution, defined as 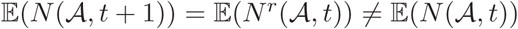 (where 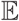 represents expectations) will occur.

In our first models we assume non-overlapping generations, 
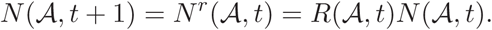
and a linear reproduction function 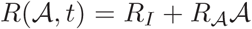 with expected fitness increasing with the value of 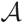. Over the course of a simulation of 30 generations (Appendix A§1.1 Model A), the population never achieves an equilibrium structure or growth rate; it grows hyper-exponentially (Figure 2(a), black line) and the shape of the breeding value distribution continually changes location (Figure 3(b), black line) and shape (Figure 2(b,d, black lines)). Linear selection only slowly erodes the genetic variance and skew (Figure 2(c,d)) and these changes lead to a slight slowing of the rate of change in the mean breeding value (Figure 2(b)) and the population growth rate (Figure 2(a)) each generation (the black lines are not linear).

**Figure 2.**
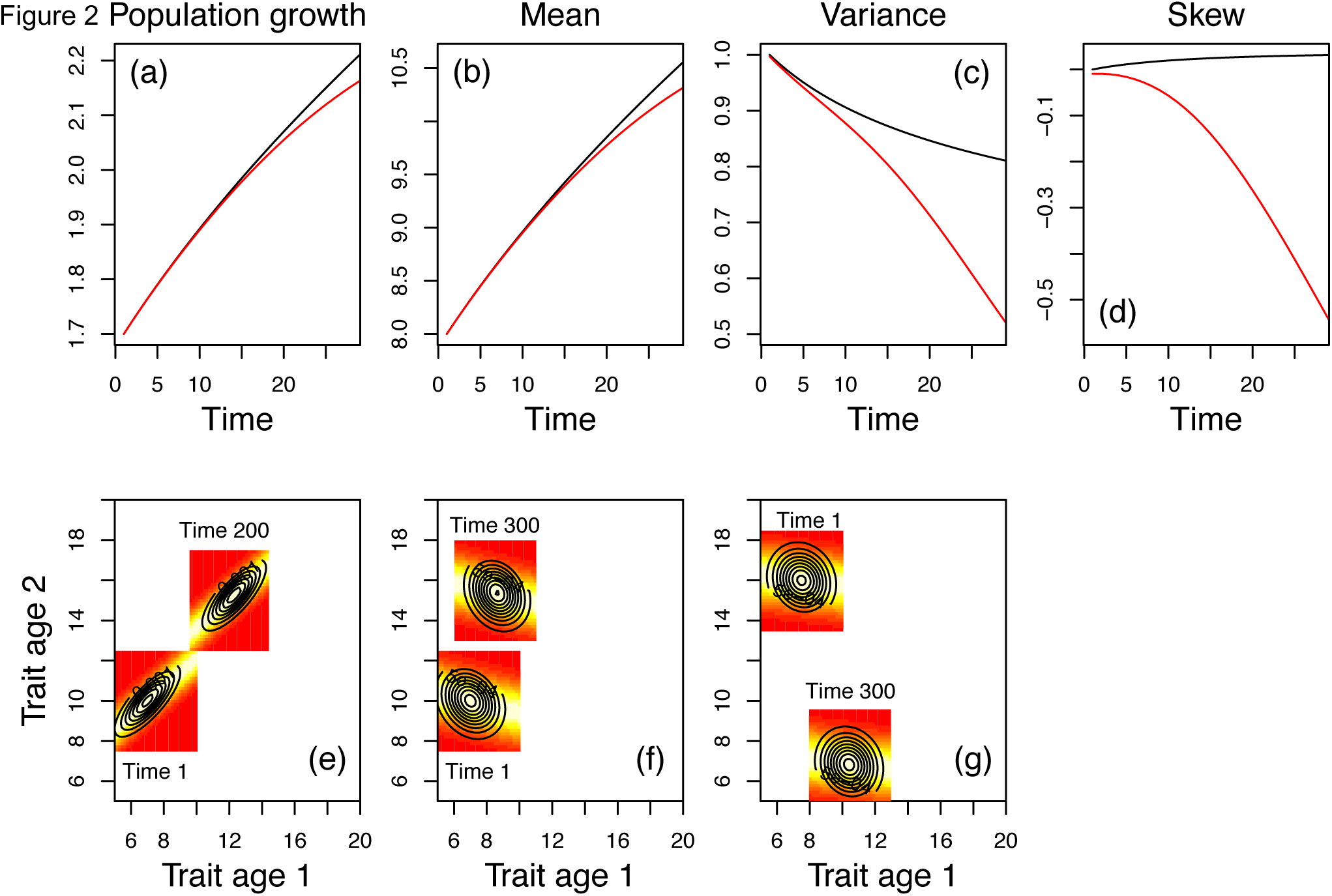
The role of selection on the dynamics of 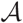. Dynamics of univariate 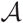 subject to linear selection and clonal inheritance (a)-(d) (Appendix A§1.1 Model A). The population does not reach an equilibrium, with (a) population growth, and the (b) mean, (c) variance and (d) skew of the character continually evolving. Imposing a maximum possible character value constrains change (red lines versus black lines (a)-(d)). In the age-structured case we track the dynamics of a bivariate character distribution (e)-(g) (Appendix A§1.1 models B, C and D). The models in (e) and (f) (Appendix AModels B and C) are identical except the starting distribution at time t = 1 has a covariance of −0.2 in (f) compared to 0.7 in (e). The parameterisation in (g) is chosen to demonstrate a case where the two traits evolve in different directions. The coloured image plots in figures (e)-(g) represent Gaussian development functions 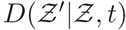 fitted to the bivariate distributions of 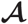 at the beginning and end of the simulation. Evolution of the bivariate character has resulted in different parameterisations of these phenomenological functions. The lighter the shading, the greater the probability of a transition from character value 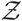 at age 1 and to 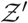 age 2.

**Figure 3.**
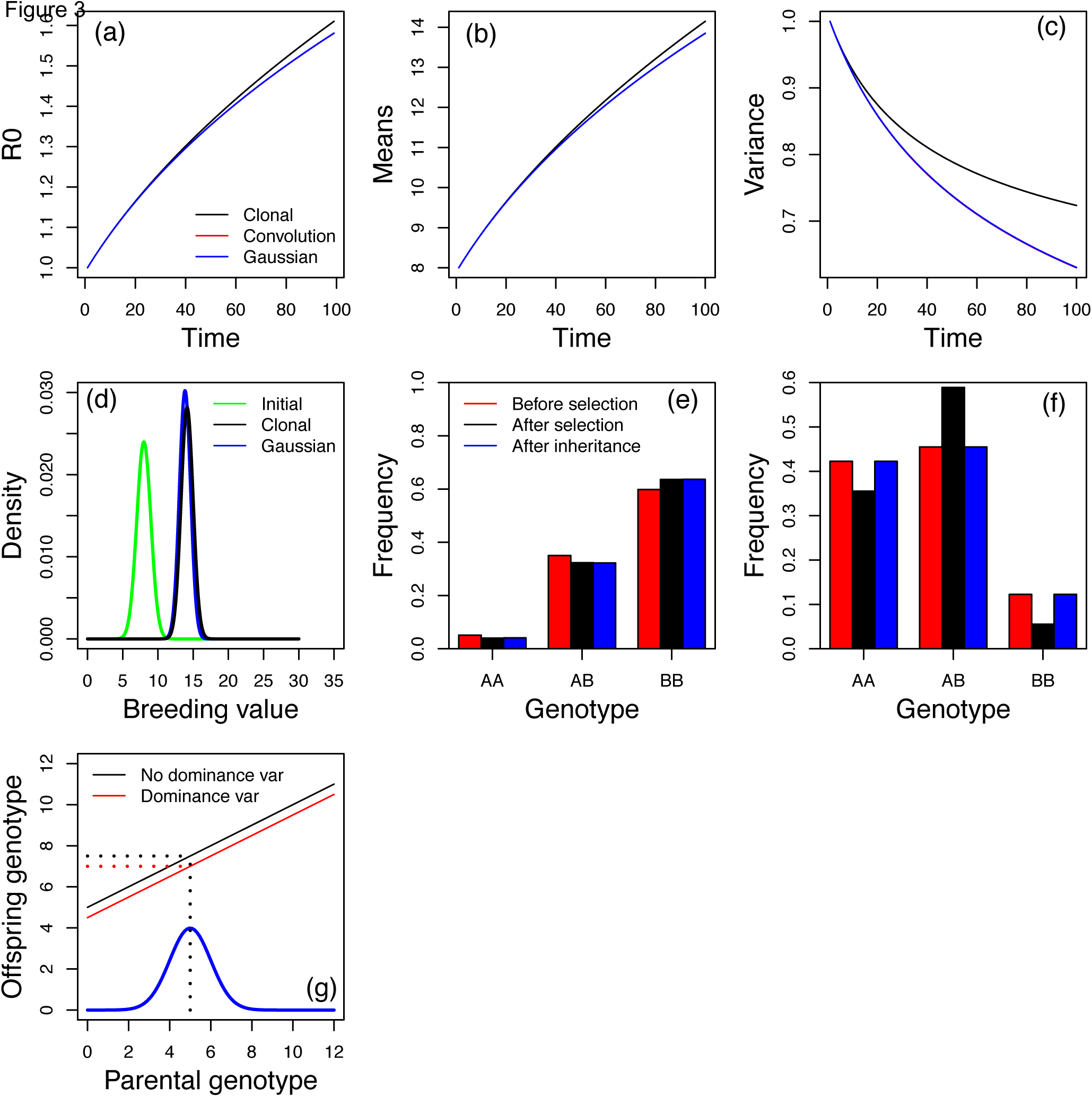
The dynamics of inheritance (Appendix AModel E). The dynamics of (a) population growth rate (R0), the (b) mean and (c) variance of 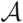 vary between models with clonal inheritance (black line), the convolution in equation (15) (red line) and the Gaussian inheritance function in equation (16) (blue line). Dynamics predicted from the convolution and the Gaussian inheritance function are indistinguishable in this model. (d) the temporal dynamics of the clonal model versus the other models. The initial distribution at t = 1 is Gaussian. After 100 generations the character distributions predicted by the clonal and sexual models have only diverged slightly. The infinitesimal model of quantitative genetics assumes that the dynamics of alleles can be inferred from the dynamics of genotypes. Under this assumption (e) selection alters genotype and allele frequencies, while inheritance does not. In contrast, (f) when dominance variance operates, both selection and inheritance alter genotype frequency while neither alter allele frequencies. For a Gaussian distributed character, (g) dominance variance acts as an offset, reducing the intercept of a Gaussian inheritance function.

In this model there are two ways to prevent the fitness function from generating change in the location of the distribution. First, the fitness function can take unimodal non-linear forms such as 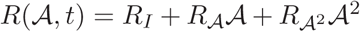 with 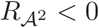 and 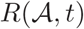 constrained to non-negative values. This generates stabilizing selection, with the mean breeding value being maintained at the value that maximizes fitness. Eventually, in this model, the breeding value distribution will achieve a trivial equilibrium – a Dirac delta function at this value. Second, continual change in the location of the distribution can be prevented by defining a maximum possible value for 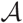 that cannot be exceeded. This captures a genetic constraint in the maximum possible character value – i.e. evolution has not evolved a genetic solution to creating a larger breeding value. In our models, this process can be captured by setting the abundance of *N*(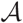 > *x*, 1) where *x* is the maximum possible trait value that evolution can achieve. Selection now pushes the breeding value distribution up to *x*, again eventually achieving a trivial equilibrium captured by a Dirac delta function where all mass of the distribution is at 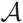 = *x*.

Genetic constraints can also impact the transient dynamics of the breeding value distribution (Figure 2(a-d, red lines)). When we impose a genetic constraint (Appendix A§1.1 model A with *x* = 11.5), the genetic variance and skew evolve faster than when no genetic constraint is in place (Figure 2(c) and (d)). These more rapid changes result in a slowing in the evolution of the mean breeding value (Figure 2(b)), and of the population growth rate (Figure 2(a)).

Genetic covariances between traits can also capture genetic constraints and can also influence the outcome of evolution. We demonstrate this by developing an age-structured model. 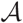 now becomes age-structured but is still inherited at birth. We construct a multivariate character 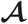 describing the breeding values that influence a character at each age (e.g. 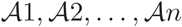 for breeding values at ages *a* = 1,2,…, *n*). If some of the same loci contribute to the genetic components of the character at different ages there is a genetic covariation across ages. The genetic variances within each age, and the covariances between ages, can be used to construct a **G** matrix (Lande, 1979). Such age-structured **G** matrices underpin the character-state approach of quantitative genetics (Lynch and Walsh, 1998). In the age-structured model that follows, we define a bivariate normal distribution with a known variance-covariance structure as our starting point and iterate this forwards (Appendix A§1.1 models B-D). We consider a simple case: a monocarpic biennial life cycle where individuals in their first year of life do not reproduce and all age 2 individuals die after reproduction. As with our model for a species with non-overlapping generations we assume clonal inheritance, 
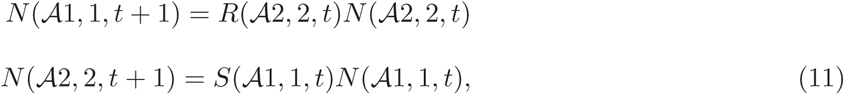
where survival from age 1 to age 2 is specified as 
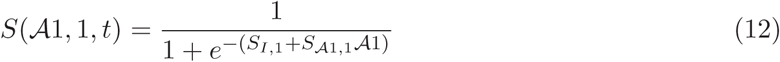
with expected survival to age 2 being highest for larger values of 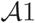. Although 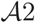 is not under direct selection, its distribution is modified by its covariance with 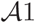.

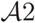, the genotype at age 2, determines expected reproduction, 
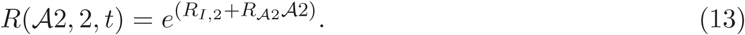

Although 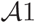 does not directly influence reproduction, there is an association between it and reproduction via its covariance with 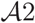. All age 2 individuals die following reproduction in this model, although it is possible to extend our approach to any arbitrary number of ages.

The evolutionary dynamics that particular parameterizations of the fitness functions 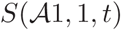 and 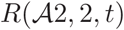 generate are dependent upon (i) the initial covariance between the characters and (ii) the fitness functions (Appendix A§1.1 models B-D). Many parameterizations and initial covariances are likely to generate evolutionary dynamics that may be biologically unrealistic. We demonstrate this with three contrasting parameterizations, considering size as our trait (Figure 2(e)-(g)). In the first example, (Figure 2(e) Appendix A §1.1 model B), the two characters positively covary and experience selection in the same direction. Over the course of the simulation the average developmental trajectory has evolved with 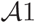 evolving to be 1.76 times larger and 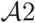 evolving to be 1.52 times larger. For a trait like body size, such a proportional change at different ages may be appropriate. In examples (Figure 2(f and g), Appendix A§1.1 models C and D) the bivariate character evolves in contrasting ways. In (F), 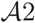 evolves much faster than 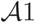 while in (G) 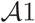 evolves to be larger, while 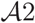 evolves to be smaller. These simulations demonstrate that only a constrained set of fitness functions and genetic covariances will give biologically realistic evolutionary trajectories for the size-related traits that biologists often study.

We now return to a univariate model and examine the clonality assumption. How can the clonality assumption be relaxed, and what are the consequences? In sexually reproducing species, offspring inherit a mix of their parent’s genomes. However, genetic segregation means that full siblings do not have the same genotype. When additivity is assumed, the breeding value of offspring is expected to be midway between parental breeding values. However, to obtain the distribution of offspring genotypes, the contribution of genetic segregation to variation among offspring needs to be taken into account. In two sex models, three steps are required to generate the distribution of offspring genotypes or breeding values given parental values. First, a distribution of mating pairs needs to be constructed. Second, the distribution of midpoint parental genotypes or breeding values given the distribution of mating pairs needs to be constructed. Third, segregation variance needs to be added to the distribution (Feldman and Cavalli-Sforza, 1979; Felsenstein, 1981; Turelli and Barton, 1994). The mating system and the segregation variance are related: when mating is assortative with respect to genotype, the segregation variance is small and siblings closely resemble one another and their parents. In contrast, when mating is disassortative with respect to genotype, siblings can differ markedly from one another, and the segregation variance is large.

Expressions have been derived for the segregation variance for the infinitesimal model where it is assumed that traits are determined by a very large number of unlinked loci of small additive effects and mating is random (Fisher, 1930). The infinitesimal model is assumed in most empirical quantitative genetic analyses (Kruuk et al., 2008) and in our initial model. For random mating where both sexes have identical demographies, the distribution of offspring breeding values given parental breeding values is (Barfield et al., 2011): 
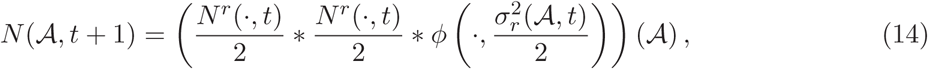
where * represents convolution and 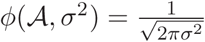 exp 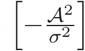 is a Gaussian function with mean zero and variance *σ*^2^ representing the segregation variance.

If males and females have different demographies then they will have different distributions of genetic values after selection; we represent these as 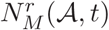 and 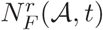, respectively. In this case, eq. (14) is replaced by 
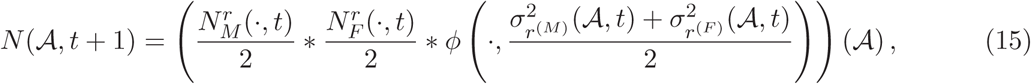
where 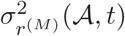 and 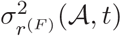 are variances of the post-recruitment-selection genetic value of males and females. respectively. We do not superscript the *rs* with *σ^2^* to avoid a notation making it appear *σ* is raised to some quantity 2*r*.

The first two terms on the right hand side of equation (15) generates the distribution of expected parental midpoint values; it ensures that the mean breeding value among offspring is midway between the two parental breeding values. However, because the parental distributions are halved, the variance of this distribution is half that of the parental distributions. The third term on the right hand side of equation (15) adds the segregation variance. For random mating, the variance is assumed to be normally distributed with a mean of 0 and a variance of half the additive genetic variance among the entire population when the population is at linkage equilibrium (Felsenstein, 1981). We approximate this variance as half the additive genetic variance in the parental distribution (Feldman and Cavalli-Sforza, 1979). This approach has already been incorporated into IPMs (Barfield et al., 2011; Childs et al., 2016).

We now run two simulations (Figure 3(a)-(d)) to examine differences in the predictions of clonal and sexual models. The first model assumes clonal inheritance and the second the convolution in Equation (15), with both models assuming a linear function 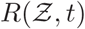 (Appendix A§1.1 model E). The two models predict slightly divergent dynamics. The reason for this is that equation (15) results in the skew and kurtosis in 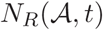 is reduced at each time step in the sexual model compared to in the clonal model. If selection is exponential (and the starting distribution proportional to a Gaussian distribution) then there will be no difference between the two approaches. This is because a normal distribution multiplied by an exponential fitness function results in a normal distribution with an unchanged variance (Diaconis et al., 1979). These results suggest that insights from clonal models will approximate those from sexual models reasonably well, at least when males and females have similar demography.

Some authors have queried the use of Equation (3) as an approximation in IPMs to the inheritance convolution in Equation (15) used in models of sexually reproducing species (Chevin et al., 2010; Janeiro et al., in press). However, being able to construct inheritance functions for 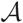 that are of the form of equation (3) would be useful as it would permit methods developed for two sex phenotypic IPMs to be applied to evolutionarily explicit IPMs (e.g. Schindler et al., 2015). Given Gaussian approximations frequently perform well in models of evolution (Turelli and Barton, 1994) we hypothesize that Gaussian inheritance functions may perform well in evolutionarily explicit IPMs. We consequently constructed a Gaussian inheritance function and compared results with those obtained from the convolution.

Equation (15) results in the mean and variance of the parental and offspring breeding value being the same. We can approximate this by ensuring that the function 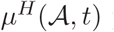 passes through the coordinate 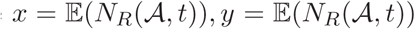 and that the variance 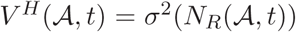. When both sexes have the same demography, we can write, 
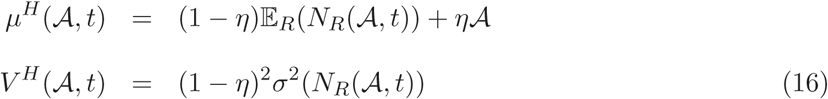
where 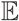 and *σ^2^* represent expectations and variances respectively and *η* represents the degree of assortative mating. When *η* mating is entirely assortative, when *θ* = 0.5 mating is random and when *η* = 0 mating is completely disassortative. An equation for the case when males and females have different demographies is provided in the Appendix A§1.3. The approximation in Equation (16) will increase in accuracy as the distribution of mid-point parental breeding values becomes more Gaussian.

When we compared predictions from equations (15) and (16) with *η* = 0.5 using the same model used to compare clonal and sexual life histories, results were indistinguishable (Figure 3(a)-(d). This reveals that, for linear selection, Gaussian inheritance functions for 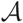 perform remarkably well.

None of our models to date include any form of mutation. We have not incorporated mutation into our models as we are simulating responses to environmental change over a few tens to hundreds of generations (Figures 1-3), and over that time period mutation is unlikely to play a major role in adaptation. However, for simulations over longer time periods, we can incorporate mutation into our models by slightly increasing the size of the segregation variance (e.g Lynch and Walsh, 1998). This will have the effect of increasing the additive genetic variance, partly countering any loss of genetic variance due to selection.

Our approximation can be used to examine the dynamical contributions of non-additive genetic processes to population responses to environmental change in a phenomenological manner. Fisher (1930) demonstrated that dominance variance can be treated as an offset, and in our models this would lower the intercept of the function 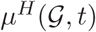 in equation (16). A consequence of this is that the mean of the offspring genotype is no longer equal to the mean of parental genotype and the dynamics of genotypes no longer exactly match the dynamics of alleles. We demonstrate this with a single locus-two allele model. When the effects of alleles are additive, the dynamics of the genotype captures the dynamics of alleles (Figure 3(e)). In contrast, when the heterozygote has higher fitness, allele frequencies do not change once the equilibrium is achieved. However, selection and inheritance alter genotype frequencies (Figure 3(f)). This effect of dominance variance can be phenomenologically capturing within an IPM by setting the intercept of the inheritance function for the genetic component of the phenotype to be less than 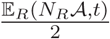 – this imposes an offset that can reverse gains made by selection (Figure 3(g)). Because this offset is negative when dominance variance is operating, dominance variance will slow, or prevent, rates of evolutionary change. We could easily phenomenologically explore how a particular value of this offset impacts predicted dynamics, however, further work is required to relate different levels of dominance variance to specific values of the offset in our models.

Having shown how IPMs can be formulated to project forwards the dynamics of the genetic component of the phenotype under a wide range of circumstances, we now turn our attention to the dynamics of the environmental component of the phenotype.

### Plasticity

Plasticity is determined by the dynamics of 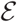 and in particular in how 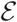 is influenced by the ecological environment *θ*. For this, we require a probability density function. We show in this section how different forms of plasticity can be incorporated into evolutionarily explicit IPMs, and explore the dynamics of some simple cases.

To capture plasticity in IPMs we need to model the probability of transition from 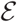 at time *t* to 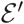 at time *t* + 1 as a function of the environment *θ*. For most plastic traits we have a poor mechanistic understanding of development and inheritance patterns, and for that reason we use the Gaussian probability density function in Equation (3).

In quantitative genetics it is often assumed that the mean of 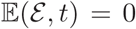 and any individual departures are purely random (Falconer, 1960). In equation 3 this requires the intercepts and slopes of the functions *μ*^D^(…) and *μ*^Η^(…) to take the following values: 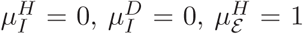 and 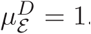. We relax this assumption and allow the mean (and variance) of 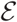 to vary with time as θ varies by specifying particular forms for development and inheritance functions of 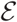.

Gaussian transition functions (equation 3) can be formulated to predictably modify moments of the distribution of 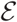 from time *t* to time *t* + 1. For example, careful choice of intercepts and slopes of 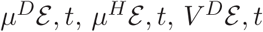 and 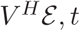 can be used to predictable grow, or shrink, the variance of 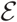 via either development or inheritance (Appendix A§1.4). In addition, specific biological processes can be easily incorporated into the dynamics of 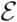: if the slopes 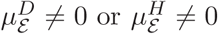 then there will be temporal autocorrelation in the value of 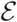 among individuals, and between parents and their offspring. For example, if 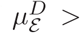 then individuals with a relatively large value of 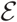 at time *t* will be expected to have a relatively large value of 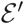 at time *t* + 1. This property of development functions is useful as it allows some memory of 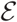 across ages: if an individual has benefited from a particularly good set of circumstances at one age, any phenotypic consequences can persist to older ages. In a similar vein, if 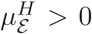 then a parent with a relatively large 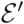 at time *t* will produce offspring with relatively large 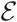 at time t + 1, a form of parental environmental effect (Nussey et al., 2007).

Different formulations of *μ*^Η^(…) and *μ^D^*(…) can be used to capture a variety of different forms of plasticity (Table 2). When *θ* is incorporated as an additive effect, it acts to shift the intercept of these functions as *t* changes. This means that the environment influences all values of 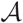 in the same manner. If 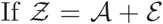 then 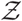 changes as a function of how *θ* influences 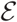 if 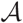 remains constant. 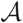 remains constant when it does not vary within individuals as they age, or if 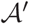 in offspring is the same as 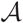 in parents.

**Table 2:**
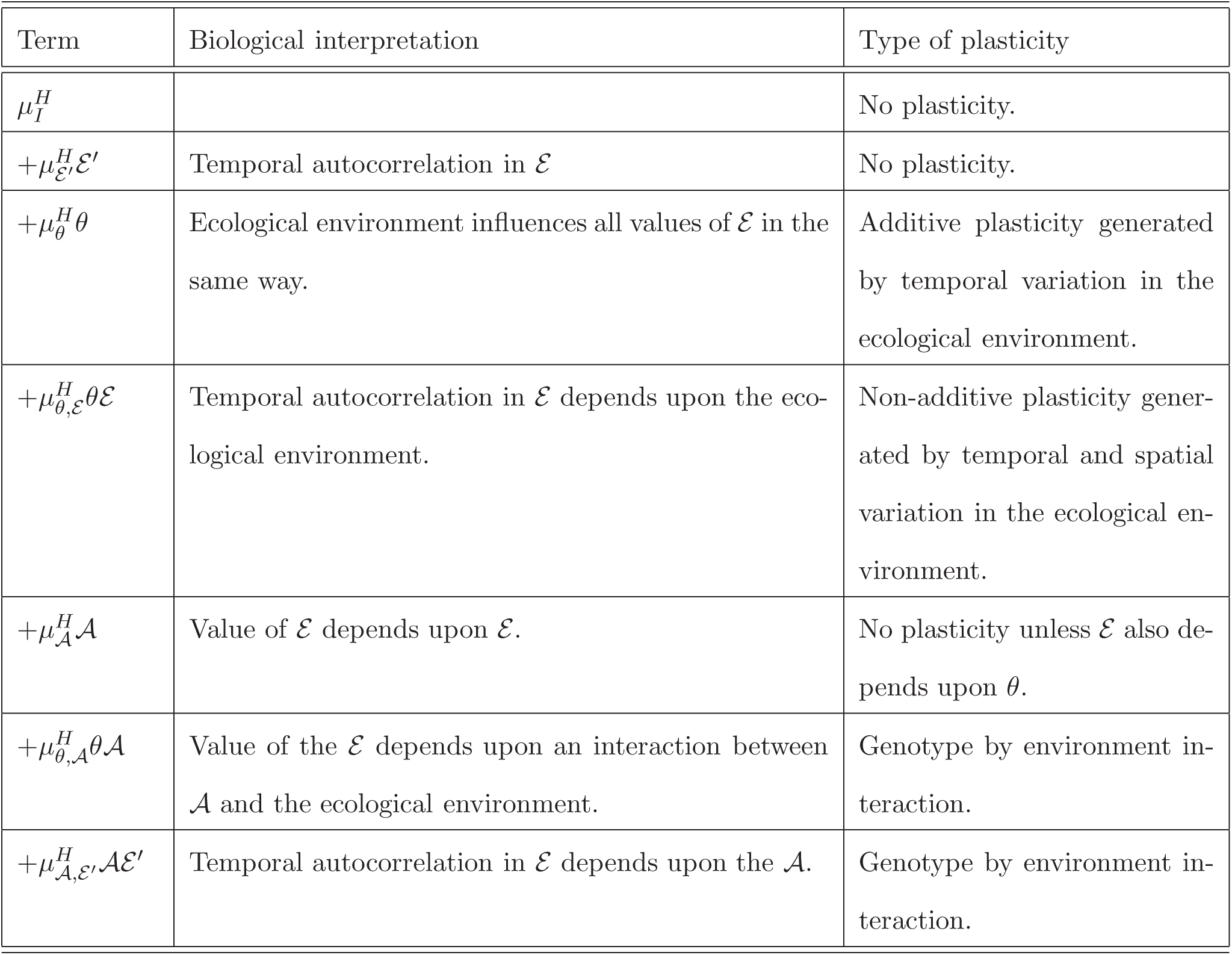
Different forms of plasticity and their incorporation into IPMs. Each term in the table below can be included in the functions 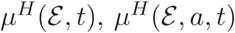 or 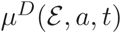. Similar terms could be included in 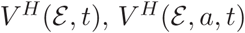 or 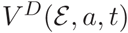 if the variance in inheritance or development varied for specific values of 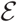 in predictable ways. This would capture different forms of bet-hedging.

Interactions between 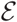,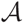 and θ are listed in Table 2. Each form describes a different type of reaction norm (Gavrilets and Scheiner, 1993). These forms allow 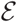 to develop among individuals (phenotypic plasticity) or be inherited (epigenetic inheritance) as a function of an individual’s breeding value 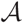 and the environment θ as well as the value of 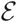 at time t.

Plasticity can be either adaptive or non-adaptive (Ghalambor et al., 2015), and both forms can be captured into our models. Adaptive plasticity enables populations to rapidly respond to an environmental change. For example, if environmental change reduces population size, then adaptive plasticity would result in a change to the mean of the phenotype via either phenotypic plasticity (the development function) or epigenetic inheritance (the inheritance function) that leads to an increase in survival or recruitment rates. In contrast, non-adaptive plasticity does the opposite, potentially exacerbating the detrimental effects of environmental change.

We demonstrate this with an example of a simple IPM of a species with non-overlapping generations: 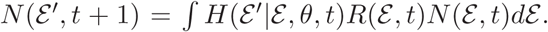. The model contains no genetic variation and the phenotype is determined by the density at the time the offspring is born. This means we can remove 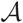 from the model. We assume a linear fitness function and a Gaussian inheritance function,

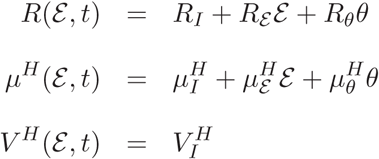

Next, we assume that the phenotypic trait is positively associated with expected recruitment such that 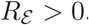. We also assume that the environmental driver is positively associated with expected recruitment such that as *θ* increases in value, fitness increases (*R_θ_* > 0). This means that the population growth rate (in a density-independent model) or population size (in a density-dependent model) also increases with *θ*. Now assume that a negative environmental perturbation decreases *θ* such that fitness decreases. For adaptive plasticity to counter this, the effect of the decrease in *θ* on epigenetic inheritance must increase the expected value of 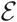. In our simple model, this can only occur if 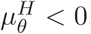 Then, as *θ* declines, 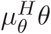 becomes less, and the value of 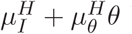 becomes larger, increasing the mean of 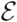 and fitness. In general, in additive linear models like this, if 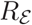 and 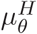 take opposing signs then plasticity will be adaptive.

We develop three density-dependent models of a phenotype in a species with non-overlapping generations. In all models we define the fitness function to be 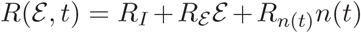 where 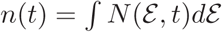 and where *R_n(t)_* < 0. In each model we define 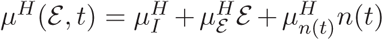. We set in model 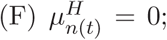 in model 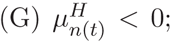 and in model 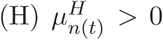 (Appendix A§1.1).

The first model (F) does not include plasticity 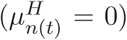 the second (G) captures adaptive plasticity 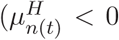 and 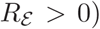, and the third (H) captures non-adaptive plasticity (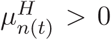) and 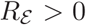. All three models include temporal autocorrelation in the environmental component of the phenotype (sometimes referred to as phenotypic carryover) when 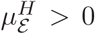 (Table 2). Because the models are not age-structured and do not include development, plasticity operates via epigenetic inheritance (e.g. maternal environmental effects). The same logic can be extended to the development function in age-structured populations. In our examples, parameterizations are chosen so all models converge to the same value of carrying capacity, *K*. Once all three models have converged, we initially impose a one off perturbation. Model (G) regains the equilibrium first, followed by model (F), and then model (H) (Figure 4(a)) showing that adaptive plasticity allows the population to recover from a one off environmental perturbation much faster than when there is no plasticity, or plasticity is non-adaptive. Non-adaptivity plasticity significantly slows the rate at which the population can recover from a perturbation, with the initial population size pre-perturbation only re-attained after 80 generations.

**Figure 4.**
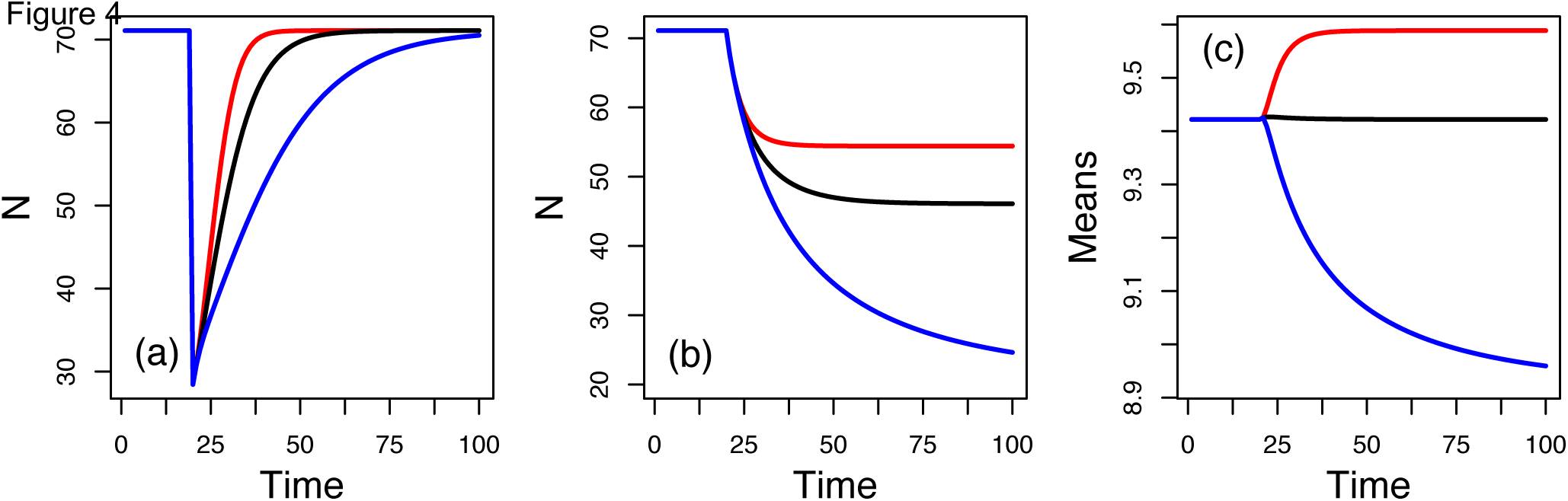
Dynamics of 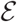 and plasticity. (a) Return times to equilibrium for three inheritance functions (Appendix A§1.1 models F-H) following a one off perturbation (see main text). There is no plasticity incorporated into model F (black line). Model G (red line) and model H (blue line) respectively incorporate adaptive and non-adaptive phenotypic plasticity. In (b) and (c) we imposed a permanent environmental change by reducing the intercept of the fitness function. (c) Represents the mean phenotype.

Adaptive and non-adaptive plasticity also impact the way populations respond to permanent environmental change. We demonstrate this by running the same models (F), (G) and (H), except now we impose a constant change in fitness by permanently changing the intercept of the fitness function *R_I_*. When we do this, the three models attain different equilibria population sizes (Figure 4(b)) and different mean phenotypes (Figure 4(c)). Model (G) achieves a larger population size than the two other models. This buffering of the population against environmental change happens because adaptive phenotypic plasticity results in a change in the mean phenotype (Figure 2(c)) that increases the expected recruitment rate and asymptotic population size (Figure 2(b)). In contrast, non-adaptive plasticity exacerbates the consequences via a change in the mean phenotype that decreases fitness.

In contrast to our example models in the §**Adaptive Evolution**, the IPMs we have developed in this section, and indeed all non-genetic IPMs so far published, achieve an asymptotic population growth rate or equilibrium population size and a stable population structure. These IPMs have monotonically increasing or decreasing fitness functions: an increase in the character results in an increase in expected fitness. A consequence of this is that in these models the recruitment function acts to alter the location of the character distribution, and often also alter its shape (Wallace et al., 2013). This is reflected in the means (and often other moments) differ between the distributions of the phenotype pre- and post-selection. In models at equilibrium with monotonic fitness functions, the inheritance function must reverse the locational and shape changes caused by the fitness function. This is because at equilibrium the moments of the phenotype distribution at times t and t + 1 must be equal.

In models of species with non-overlapping generations at equilibrium like those above, the inheritance function for 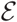 must exactly reverse the changes to the character distribution generated by the fitness function. This requires moments of parental and offspring characters to differ from one another if 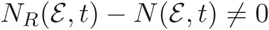. When there is a correlation between parental and offspring traits in the inheritance function for 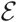 as in our models, the intercept of the inheritance function must take a value such that offspring characters are smaller than their parent’s were at the same age (Coulson and Tuljapurkar, 2008).

IPMs for species with overlapping generations include development functions 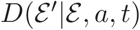. These functions can alter the size (population size) and shape of the distribution of 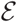 as individuals age. When generations are overlapping, and at equilibrium, changes to the location of the character distribution via survival, recruitment and development are all exactly countered by the inheritance functions 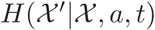.

Coulson and Tuljapurkar (2008) showed that in red deer age-specific effects meant that young and old parents were incapable of producing offspring that had the same body weight as they did at birth. This process reversed the effects of viability selection removing small individuals from the population early in life. The same process was observed in marmots (Ozgul et al., 2010) and Soay sheep (Ozgul et al., 2009) and may be general for body size in mammals.

The models we have developed do not incorporate the evolution of phenotypic plasticity. However, if genotype-by-environment interactions were included in models, such that different breeding values had different responses to environmental variation, then plasticity could evolve. If this was coupled with a segregation variance that introduced novel genetic variance, this could capture the evolution of novel phenotypic plasticity. However, over the time periods over which our simulations are conducted, the evolution of novel forms of phenotypic plasticity, is unlikely to play a major role in population responses to environmental change.

We have now developed IPMs for (i) 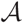 where we assumed all individuals had the same, constant, 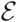 and (ii) 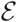 where we assumed all individuals had the same, constant, 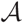. We have shown how IPMs can capture a wide range of biological processes including adaptive and non-adaptive plasticity and correlated characters, and the circumstances when equilibria are achieved. We now link together these advances into models of the joint dynamics of the bivariate distribution 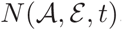.

### Models for the phenotype consisting of genetic and environmental components

In the section we construct models where the character can be determined by a mixture of the genetic and environmental components. These models allow us to explore how adaptive evolution is influenced by plasticity.

We first develop a dynamic univariate version of the Breeders equation (Falconer, 1960) for a species with non-overlapping generations in a constant environment. In this case, the environmental component of the phenotype is assumed to be a consequence of developmental noise: individuals achieve their genetic potential, plus or minus a departure. At each generation within each breeding value, the distribution of the environmental component of the phenotype is assumed to be Gaussian with a mean of 0 and a constant variance (Appendix A§1.1 Model I).

Our initial conditions are a bivariate Gaussian distribution of 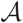 and 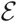 which we iterate forwards for 300 time steps. Over time, the mean of the genetic component of the phenotype increases. In contrast, the mean of the environmental component is constant. The population grows hyper-exponentially (Figure 5(a)), the mean of the phenotype increases in value due to evolution (Figure 5(a,d)) and the additive genetic variance is slowly eroded (Figure A2). Because the additive genetic variance is eroded, while the phenotypic variance remains constant, the heritability declines over time (Figure A2).

**Figure 5.**
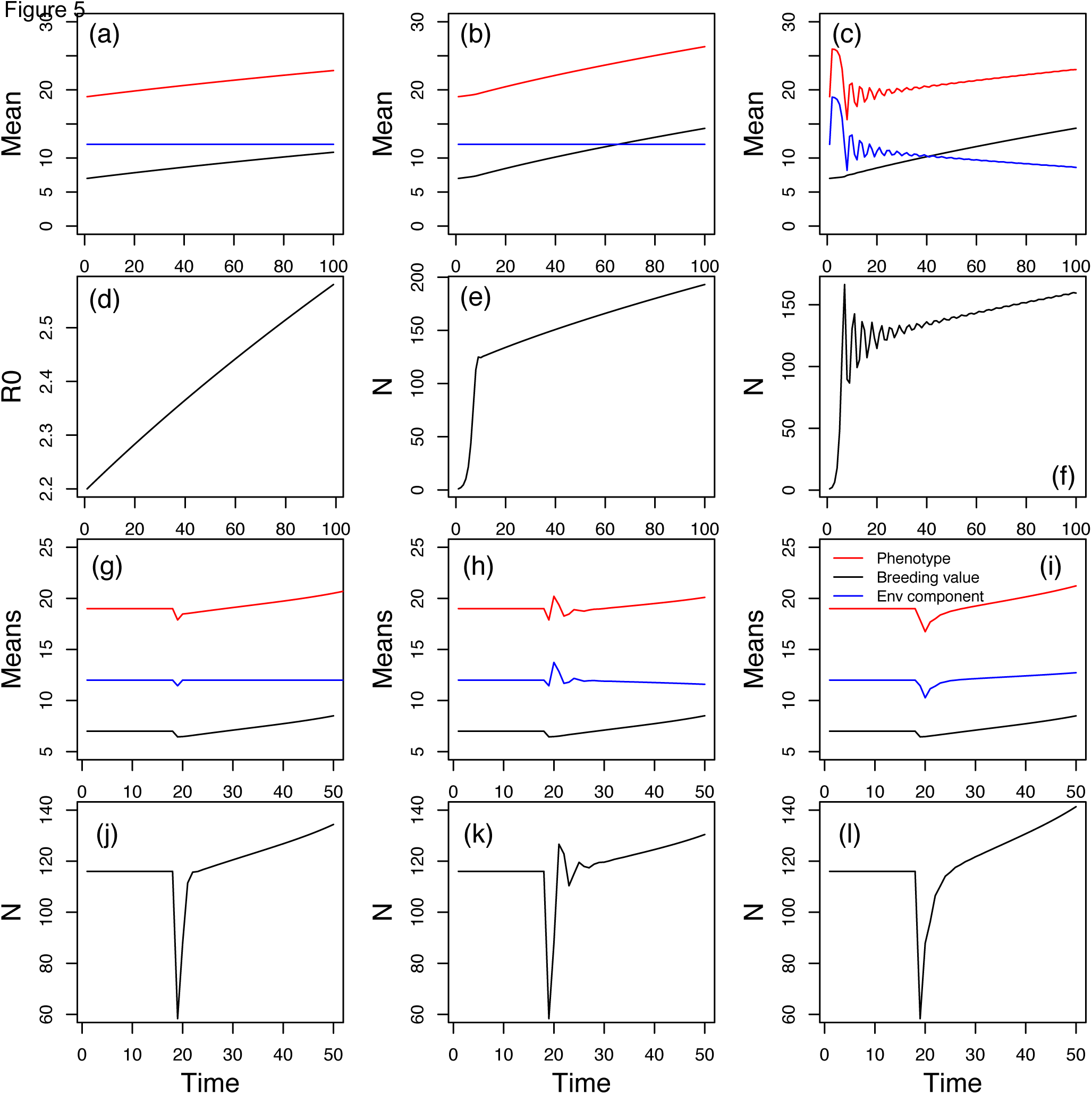
A dynamic version of the Breeders Equation. The dynamics of the phenotype (red lines) and its genetic (black lines) and environmental (blue lines) components (a)-(c) and (g)-(i), and the dynamics of the population (d)-(f) and (j)-(l). In the first model (a) and (d), both fitness and inheritance of the environmental component of the phenotype are independent of density (Appendix A§1.1 model I). In the second model (b) and (e) fitness is negatively density-dependent and inheritance of the environmental component of the phenotype is density-independent (Appendix A§1.1 model J). In the third model, both fitness and inheritance of the environmental component of the phenotype are negative density-dependent (Appendix A§1.1 Model K). The treatment of plasticity can dramatically influence the dynamics of the phenotype and population size (Appendix A§1.1 models L-N). Adaptive phenotypic plasticity (h) and (k) leads to the population size and phenotype recovering from a perturbation much faster than non-adaptive plasticity (i)-(l). The absence of a plastic response (g) and (j) results in the population recovering from a perturbation at an intermediate rate between cases where adaptive and non-adaptive plasticity are operating.

Our second model (Appendix A§1.1 model J) has a negative density-dependent term in the fitness function. The phenotype evolves faster in this model than in our density-independent model (Figure 5(b)). Population size grows nearly linearly in this model (Figure 5(d)), although the rate of increase does slow slightly each generation as genetic variation is eroded. The difference between the hyper-exponential (density-independent model) and nearly linear increases (density-dependent model) in population size explain the difference in the rates of evolution. This is because the selection differential that determines the rate of evolution (an emergent property from our model (Wallace et al., 2013)) has the population growth rate in its denominator. The population growth rate is smaller in the density-dependent model (just above unity) than in our density-independent one (it increases with time), and this leads to an increase in the strength of selection and the rate of evolution (see also Pelletier and Coulson, 2012). A consequence of this is that the additive genetic variation and heritability tend towards zero faster the in density-dependent model than in the density-independent one (Figure A2).

In our third model (Appendix A§1.1 model K), negative density-dependence is included in the inheritance function for the environmental component of the phenotype as well as in the fitness function. This captures adaptive phenotypic plasticity. This results in a negative change in the mean of the environmental component of the phenotype with time (Figure 5(c)). This decrease is reflected in a change in the mean of the phenotype itself. Adaptive phenotypic plasticity leads to a decline in the population growth rate which results in a slight increase in the rate of evolution compared to the density-dependent model with no plasticity. However, the effect is not large and is only just distinguishable when comparing Figures 5(b) and (c).

In our final models (Appendix A§1.1 models L to N) we examine how a one off perturbation influences the mean of the phenotype, its components and the population growth rate (Figure 5(g)-(l)) when there is no plasticity, adaptive plasticity and non-adaptive plasticity. We set the variance in the genetic and environmental component of the phenotype to be equal, giving an initial heritability of *h^2^* = 0.5. In each model we allow the population to achieve the same equilibrium population size in the absence of selection 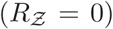. We then impose a one off mortality event when 99% of individuals above the mean of the phenotype are killed off. At this point we also impose selection 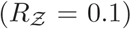. In all three models the mortality event results in a small change in the mean value of the phenotype (Appendix A§1.5 for an explanation) (Figure 5(g)-(i), red lines) but a halving of population size (Figure 5(j)-(l)). Adaptive plasticity results in the environmental component of the phenotype returning to its pre-perturbation value very quickly (Figure 5(g)-(i) blue lines). In contrast, although the perturbation causes a modest change in the mean of the genetic component of the phenotype, it takes > 10 generations for evolution to reverse the change (Figure 5(g)-(i), black lines). This demonstrates that a strong selective effect can leave a large population dynamic impact, but leave only a small initial signature in the phenotype even when the trait is highly heritable.

Over the longer term, the dynamics of the all components of the phenotype, the phenotype itself and the population dynamics all depend upon whether plasticity is adaptive or non-adaptive. Adaptive plasticity allows the population size to initially recover from the perturbation more quickly than when plasticity is absent or non-adaptive (Figure 5(j)-(l)). However, over a longer time period, non-adaptive plasticity results in the population achieving a larger size than when plasticity is absent or adaptive. These differences in population growth rate impact rates of evolution: immediately following the perturbation, the rate of evolution is greatest when plasticity is non-adaptive. However, the rate of evolution then increases when plasticity is adaptive (Figures S2 and S3). As with our previous models, the effects of adaptive and non-adaptive plasticity on rates of evolution are relatively small, but our results demonstrate how the two processes can interact.

### Signatures of evolution in models that are not evolutionarily explicit

The models in the previous section are quite complex. Do we always need to construct such evolutionarily explicit IPMs to predict population responses to environmental change, or can we rely on simpler, phenotypic IPMs? There are two reasons why it may be preferable to not construct evolutionarily explicit models. First, evolutionarily explicit IPMs are more complicated to construct than those that do not include genotypes or breeding values. Second, when data are unavailable to explicitly include breeding values into models (Traill et al., 2014*b*), the effects of evolution on predicted dynamics can still be explored by examining the consequences of perturbing parameter values (Traill et al., 2014a).

When evolution occurs within a system we would expect parameters in phenomenological inheritance and development functions that are fitted to data to change with time. We can see this in Figure 2(e)-(g)). In these age-structured evolutionarily explicit models, the bivariate breeding value distribution (black contours) changes location as evolution occurs. We have fitted Gaussian development functions to these bivariate distributions at the beginning of each simulation and at the end (coloured image plots). The parameters that determine these developments functions have clearly changed as the location of the functions have changed. A similar process occurs for inheritance functions (not shown).

Numerous authors have previously noted this phenomenon in models of evolution. For example, in population genetic (Charlesworth, 1994) and eco-evolutionary models (Coulson et al., 2011; Yoshida et al., 2003) when genotype frequencies change with time, macroscopic, population level quantities like mean survival and recruitment also change; in adaptive dynamic models, as one strategy invades another, population level parameters inevitably change with strategy frequency over time (Metz et al., 1996); in quantitative genetic predator-prey models population level parameters of both predators and prey vary over time leading to persistence of the interaction (Doebeli, 1997); and in evolutionarily explicit IPMs parameters in inheritance functions have been shown to change with time as evolution progresses (Rees and Ellner, 2016). These insights are useful because if evolution is occurring within a system, then temporal trends in statistical estimates of model parameters would be expected – in other words, the effect of time, either additively or in an interaction with other parameters, would be expected in 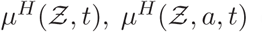 or 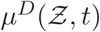. If marked temporal trends are observed in parameters in development and inheritance functions that cannot be attributed to a changing environmental driver, then evolutionarily explicit IPMs may be required.

What about parameters in fitness functions 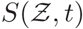 and 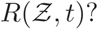 Can any inferences from temporal trends in these parameters be made? In our approach, evolution of a focal trait would not be expected to alter statistical estimates of the fitness functions. In our models, evolution simply moves the location and shape of the phenotype distribution, but not its association with survival or recruitment.

We have identified one circumstance where evolution will leave a signature in the dynamics of fitness function parameters. Parameters in these functions can evolve in the presence of a genetically unmeasured correlated character that is also evolving. To demonstrate this we construct a model of a bivariate character, examine the dynamics it predicts, before exploring the consequences of failing to measure one of the characters.

We assume clonal inheritance such that dynamics of the characters are solely determined by a bivariate fitness function,

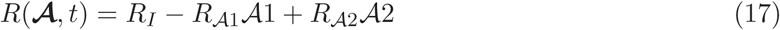

The dynamics this model predicts depend upon the initial covariance between the two characters in a similar way to our age-structured model (equation 11). In our first example the two characters negatively covary, while in the second they positively covary (Appendix A§1.1 for model parameterizations). The initial negative covariation allows rapid evolution, with population growth (Figure 6(a)), the mean of the characters (Figure 6(b)), their variances (Figure 6(c))) and the covariance between them (Figure 6(d)) evolving relatively quickly. In contrast, when the two characters positively covary, evolution is much slower, with the character means, variances and covariance changing much more slowly, even though the fitness functions are identical in each model (Figure 6(e)-(h)).

**Figure 6.**
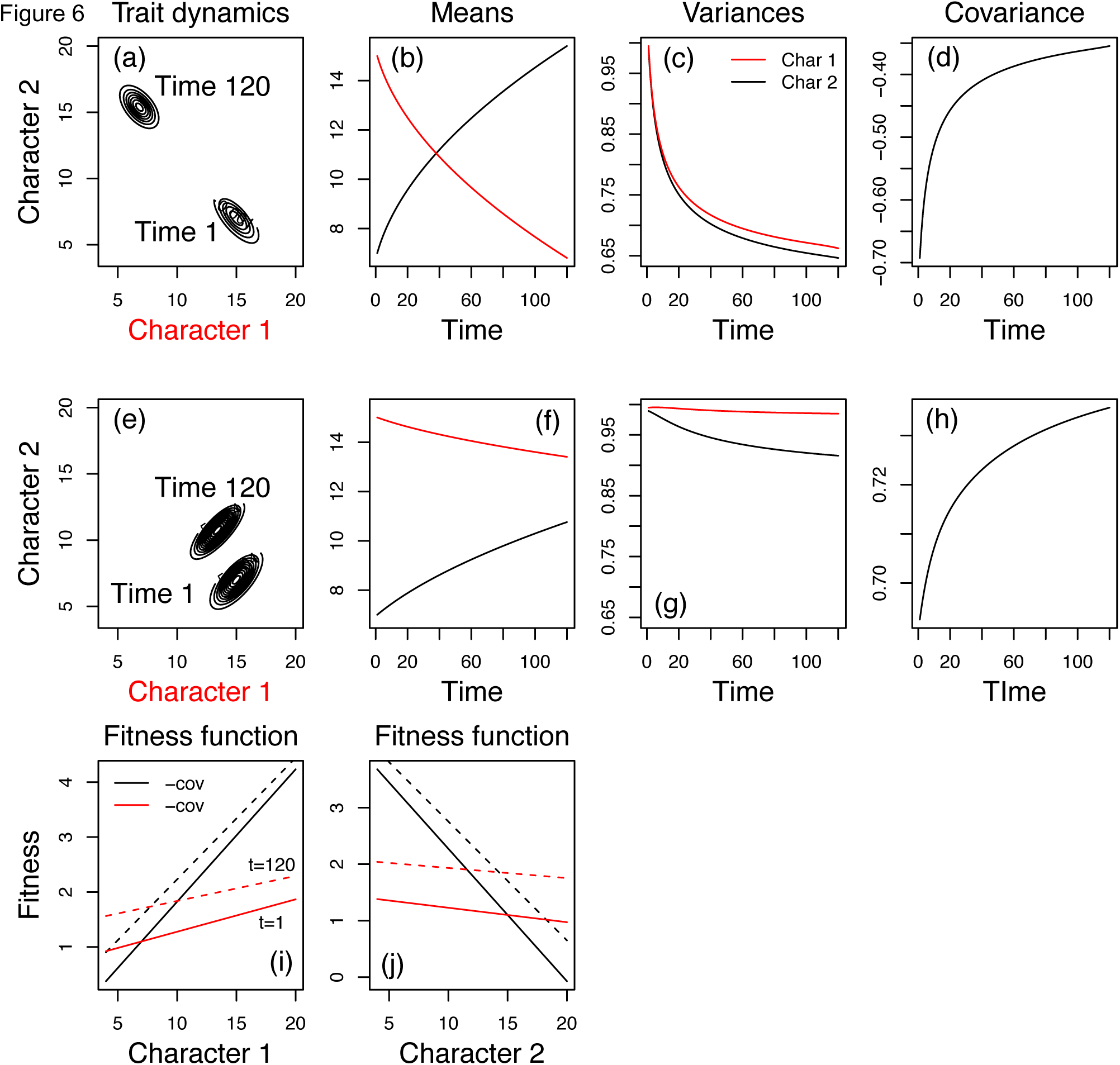
Dynamics of bivariate characters and evolution of fitness functions in the presence of an unmeasured, genetically correlated character (Appendix A§1.1 model P and Q). We construct a simple model with clonal inheritance of two correlated characters that both influence fitness. We explore two initial starting conditions that only differ in their genetic covariance (Appendix A§1.1 models P and Q). In (a)-(d) the covariance accelerates the rate of evolution compared to (e)-(h). The dynamics of the fitness function for each character when the other character is not measured (i) and (j). Regardless of the covariance between characters, the fitness functions evolve during the course of 120 time step simulation.

We now construct a fitness function for 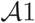 when 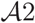 is not measured. We start by defining mean fitness, an observable, as 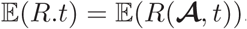. The slope 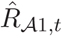 is given by, 
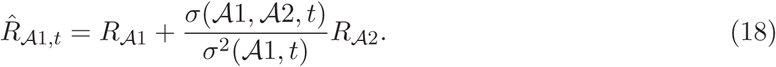

The intercept can be calculated in the usual manner by estimating the means of fitness and 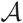 
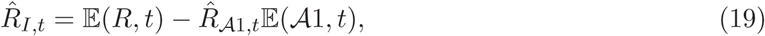
giving, 
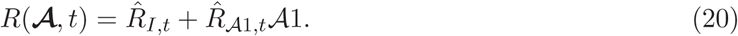

Equation (20) is what would be estimated from data if 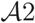 were not measured and included in analyses (Kendall, 2015; ?). It will correctly describe the consequences of selection on 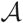 even though 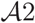 could be correlated with it. This is because the unmeasured correlated character impacts fitness whether it is measured or not, and consequently impacts the association between the focal character and fitness in its absence (?). However, the fitness function cannot provide accurate predictions over multiple generations when it is assumed that the fitness function is constant.

Over multiple generations the existence of unmeasured correlated characters will alter parameters in the fitness function in Equation (20) if selection alters genetic variances and covariances of measured and unmeasured correlated characters (Figure 6(i)-(j)). This is because 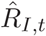 and 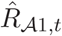 are both functions of the covariance between the two characters (equations 18-20). If selection alters this covariance, parameters 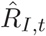 and 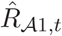 will evolve with time. It is also why we use the subscript *t* for 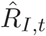 and 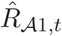. Evidence of correlated characters under selection can consequently be inferred if parameters in fitness functions are observed to change with time in a system in the absence of a changing environmental driver. Note that a non-stationary unmeasured environmental driver could also generate trends in parameter values in fitness functions in phenomenological IPMs.

## Discussion

In this paper we develop an approach that allows prediction of how populations respond to environmental change via adaptive evolution and plasticity. We do this by incorporating insights from evolutionary genetics into data-driven structured population models. Our approach is to split the phenotype into its genetic and environmental components and to model the dynamics of the genetic component with functions based on understanding of the mechanisms of inheritance. In contrast, the dynamics of the environmental component of the phenotype are modeled with phenomenological functions that can be identified from the analysis of data. Our approach is appropriate for sexually reproducing or clonal species with either overlapping or non-overlapping generations.

### Evolutionarily explicit structured models

IPMs are now a widely used tool in ecology and evolution because of their versatility and the ease with which they can be parameterized (Merow et al., 2014). All key statistics routinely estimated in population ecology, quantitative genetics, population genetics and life history describe some aspect of a character distribution or its dynamics (Coulson et al., 2010). IPMs are so versatile because they describe the dynamics of these distributions. Characterization of the determinants of these statistics gained via sensitivity or elasticity analysis of models have provided insight into how ecological and evolutionary quantities that interest biologists are linked (Coulson et al., 2011). Although this logic was developed several years ago, there has recently been criticism that IPMs cannot be used to track the dynamics of multivariate breeding values expressed at different ages (Chevin, 2015; Janeiro et al., in press). Our paper addresses this criticism head-on—we show how IPMs can be formulated to capture a mechanistic understanding of inheritance and development. In demonstrating this we develop a general modeling approach to capture population responses to environmental change. Having done this, we are now in a position to construct IPMs of quantitative characters and examine how perturbing the environment will influence not only the dynamics of the phenotype and its genetic and environmental components, but also the life history (Steiner et al., 2014, 2012) and population dynamics (Easterling et al., 2000).

The work we present here adds to a growing literature that explicitly incorporates evolution into structured models, and IPMs in particular. Within the population genetics paradigm, Charlesworth (1994) developed structured models with a one-to-one map between genotype and phenotype in age-structured populations. Building on this work, Coulson et al. (2011) showed how simple genetic architectures can be incorporated into IPMs, developing a model to explore how evolution at a single locus would occur simultaneously with phenotypic change of discrete and continuous characters, life history and population dynamics.

Working in the quantitative genetic paradigm, Lande (1982) derived age-structured models that tracked the dynamics of the mean of the additive genetic component of the phenotype (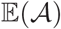 in our notation) and the mean of the phenotype itself 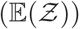. He assumed a constant genetic-variance covariance matrix and consequently weak selection and normally distributed character values—assumptions we relax. Barfield et al. (2011) extended Lande (1982)’s approach to track the dynamics of the entire character distribution and to stage-structured populations. In doing so, they developed a general, flexible approach to track the entire distributions of 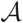 and 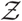. Childs et al. (2016) extended this approach to two sexes. Because 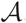 is inherited with mechanistic rules that are not impacted by the environment, while inheritance and development of [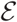 are plastic and can be impacted by the ecological environment (Falconer, 1960), it is difficult to incorporate the effects of the environment on the dynamics of the phenotype by focusing on 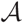 and 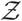 as Lande (1982), Barfield et al. (2011) and Childs et al. (2016) have done. In contrast, our approach (which otherwise has a similar logic to Barfield et al. (2011) and Childs et al. (2016)) tracks the dynamics of 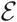 and 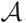 (or 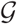—the full genotypic value, including non-additive components—if desired), making incorporation of environmental drivers that influence inheritance and development of [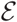] more straightforward. We show that it is possible to have selection operating on the phenotype while incorporating modern understanding of genetic inheritance into the dynamics of the genetic component of the phenotype and phenomenological insight into the role of the ecological environment on the dynamics of the environmental component of the phenotype. By doing this, we show how population responses to environmental change via adaptive evolution, phenotypic plasticity and epigenetic inheritance can be simultaneously explored. This opens up the way to provide novel insights into the circumstances when each process is expected to contribute to population responses to environmental change.

### Population responses to environmental change

Unlike previous evolutionarily explicit IPMs (Barfield et al., 2011; Childs et al., 2016; Rees and Ellner, 2016), our approach requires explicit consideration of the inheritance and development of 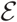, the environmental component of the phenotype. This allows our models to capture a range of plastic responses to environmental change along with adaptive ones. What do our findings say about the contributions of plasticity, evolution, and their interaction to population responses to environmental change?

Detrimental environmental change often causes a decline in population size. When there is an association between a phenotypic trait and survival and recruitment rates, phenotypic change can lead to increased survival and recruitment rates (Ozgul et al., 2010) and consequently an increase in population growth rate and size. Two processes can lead to phenotypic change - plasticity and adaptive evolution. There has been considerable discussion about the relative roles of each in allowing populations to respond to change (e.g. Bonduriansky et al., 2012; Chevin et al., 2010).

Genotypes and breeding values remain fixed within individuals throughout life which means that differential survival and recruitment rates are the processes that alter these distributions and underpin evolution. The strength of differential survival and recruitment can be impacted by environmental variation generating fluctuating selection (?). Environmental variation does not influence genetic inheritance: once mating pairs are formed, inheritance of breeding values, 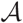, does not alter the mean or variance of breeding value distributions (Fisher, 1930). In contrast, distributions of the environmental component of the phenotype can be altered via survival, recruitment, development and inheritance with each process potentially impacted by environmental variation (Reed et al., 2010). Given these differences between the dynamics of 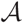 and 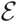 plasticity can lead to more rapid change than evolution in our models (e.g. Figure 5). This is because more biological processes can directly alter the distribution of plastic characters than can impact distributions of breeding values. These results are consistent with those of other authors, including Lande (2009) and Chevin et al. (2010), who also concluded that plastic change should be faster than evolutionary change. But how quickly will evolution alter phenotypic trait distributions?

Our results on the speed of evolution suggest that claims of detectable rapid evolution in quantitative phenotypes is likely to take a few tens of generations. For example, environmental change increases mortality leading to a decline in population size, but for mortality selection to lead to evolutionary change over the course of a generation, a large proportion of the population needs to be selectively removed and the phenotype needs to be highly heritable. This is seen in our model results (Figure 5(g)-(i)) and with a simple numerical example: when all individuals above the mean of a normally distributed character are removed from the population and the trait has a heritable of *h^2^* = 0.5, population size halves in a single time step but the mean of the character will only shift from the 50^th^ percentile to the 37.5^th^ percentile. For a standard normal distribution with a mean of 0 and a standard deviation of unity, this means the mean would only shift by 0.319 – i.e. less than 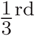 of a standard deviation – i.e. a long way from statistical significance. In reality, mortality selection resulting from environmental change will likely result in a change to the mean of the distribution that is only a fraction of a standard deviation compared to our example. Given this, reports of rapid evolution due to environmental change increasing mortality selection over a small number of generations (e.g. Coltman et al., 2003) should be treated with extreme caution. It is much more likely that change is a consequence of phenotypic plasticity. Over multiple generations, recruitment selection can also contribute to evolutionary change and our approach allows the role of this to be investigated. However, unless reproduction is restricted to individuals with extreme phenotypic trait values in both sexes, it seems unlikely that evolution can generate statistically demonstrable evolutionary change over a small number of generations (?). This is not to say that evolution is not important over longer time scales. Over tens of generations evolution can shift phenotypic trait means to a greater extent than phenotypic plasticity (Figure 5(g)-(i) blue versus black lines).

In order for plasticity to allow populations to rapidly respond to environmental change, a large proportion of individuals within the population must exhibit the same plastic response. A good example of such a dynamic is for size-related traits that are determined by resource availability, particularly when scramble competition is operating. When resources becoming limiting, all individuals will be unable to develop as rapidly as when resources are more common. A consequence of this is that individuals that developed in cohorts when resource were sparse will exhibit smaller body sizes compared to individuals in those cohorts that developed when resources were more abundant. We can capture this form of plasticity in our framework with an additive effect of density in the inheritance or development function for 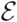 (e.g. Figure 4). In contrast, when contest competition operates, larger individuals would acquire more resources than those that are smaller, and would develop faster. We can capture this in our models with interactions between density, 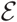 and 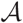 in either the inheritance or development functions for 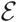.

The above discussion demonstrates how our approach can be used to capture different forms of plasticity. However, for plasticity to help populations respond to environmental change it must be adaptive: plasticity must change the mean trait value in a way that increases fitness (Ghalambor et al., 2007). We demonstrate that for additive, linear models, adaptive and non-adaptive plasticity can be specified by altering the sign for the effect of the environment in the function specifying the mean dynamics of the inheritance or development functions (Figure 4). When interactions are included in these functions specifying general rules for whether plasticity is adaptive or non-adaptive will likely be more challenging. However, our approach provides a way in which to investigate when plasticity is adaptive or non-adaptive, and how different types of plasticity will influence population responses to environmental change.

Our results also show how plasticity can influence evolutionary rates. Plasticity, operating via development and inheritance functions for the environmental component of the phenotype, alters the distribution of the phenotype, and this can alter the strength of selection, which can then influence the dynamics of the genetic component of the phenotype (evolution). The effects of plasticity on selection and evolution can be surprisingly complex. We only examined the evolutionary consequences of plasticity following an environmental shock that influenced all individuals in the same way, but even in this simple case we found that adaptive plasticity initially slowed the rate of evolution compared to non-adaptive plasticity, before increasing it (Figure 5 and Appendix A). In general in order to understand how plasticity will influence selection, it is necessary to understand how it influences both the numerator and denominator of the selection differential that underpins evolution (Pelletier and Coulson, 2012). The numerator is the covariance between the phenotype and absolute fitness (Falconer, 1960) and the denominator is mean fitness. In our models of species with non-overlapping generations this is mean recruitment – the population growth rate (Fisher, 1930). Selection is linear in our models where plasticity influences all individuals in the same way via an additive effect of density on inheritance of the environmental component of the phenotype (figure 5), and this means that plasticity influences the population growth rate rather than the numerator of the selection differential. A consequence of this is that it is differences in the population growth rate that generates the differences in evolutionary rates between models when plasticity is adaptive and non-adaptive. In more complex cases when plasticity influences the covariance between the phenotype and fitness via genotype-phenotype interactions within a generation, to understand how selection influences evolution it is necessary to understand how plasticity not only influences mean fitness, but also how it generates differences between the covariance between the genetic component of the phenotype and fitness and the covariance between the phenotype itself and fitness. Because the components of the selection differential can be calculated from IPMs (Coulson et al., 2010; Wallace et al., 2013) the approach we develop here provides a flexible way to examine how different types of plasticity can influence evolution following environmental change.

We have not considered bet-hedging in this paper. Bet-hedging is another form of plasticity that can influence the way populations respond to environmental change and it can be incorporated into IPMs (?). Deterministic IPMs incorporate probabilistic transitions when 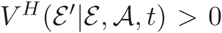 and 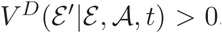. These probabilities do not vary from one time step to the next. In stochastic models these functions can include terms for an environmental driver *θ*, such that the variation in trajectories changes with the environment. In evolutionarily explicit models, the variance in transition rates among different values of 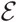 can be made to depend upon *θ*, 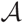 and their interaction (if desired). This means that individuals with specific values of 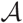 can produce offspring with more variable values of 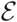 (and consequently 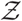) in particular environments than individuals with other values of 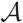. In this paper we focused on the incorporation of *θ* into 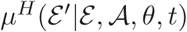 and 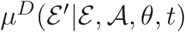 but responses to environmental change could also be incorporated into functions for the standard deviation that we use to construct our kernels.

In order to explore how the various forms of plasticity influence rates of evolution for real systems it will be necessary to parameterize our models with data.

### Parameterizing and analyzing evolutionarily explicit IPMs

A large literature exists on how to statistically parameterize IPMs (Easterling et al., 2000; Merow et al., 2014; Rees et al., 2014). The vast majority of IPMs have been constructed phenomenologically, using statistical descriptions of observational data. Several authors have shown how fixed and random effects incorporated into these statistical functions can be formulated within IPMs (Childs et al., 2003*a*; Coulson, 2012; Rees and Ellner, 2009), but additional statistical estimation is required to parameterize the evolutionarily explicit IPMs we have developed.

Fitness functions in evolutionarily explicit IPMs can be parameterized using standard general, generalized and additive regression methods that are routinely used to parameterize phenomenological IPMs (Rees and Ellner, 2009). If relatedness information is available and the infinitesimal model is assumed, genetic and phenotypic variances and covariances can be estimated using the animal model (Lynch and Walsh, 1998). These quantities can be used to construct the initial distributions of the genetic and environmental components of the phenotype. Parameter estimates of ecological drivers fitted as fixed or random effects in the animal model can be used to parameterize inheritance and development functions for the environmental component of the phenotype. It is consequently possible to parameterize models using our approach with existing methods.

There is also a large literature on how to analyze IPMs (Ellner and Rees, 2006; Steiner et al.,2014, 2012). The majority of these tools, including sensitivity and elasticity analysis of model predictions to transition rates and function parameters (Coulson et al., 2011, 2010; Ellner and Rees, 2006; Steiner et al., 2014, 2012), are likely sufficiently general to be applicable to evolutionarily explicit IPMs. In future work we plan to parameterize models for bird, mammal and fish species with overlapping generations and to analyze them with existing methods. Once evolutionarily explicit IPMs have been parameterized and analyzed we will be able to explore how populations, phenotypic characters and life histories are predicted to respond to a range of environmental changes via plasticity and adaptation.

### When should evolutionarily explicit IPMs be used to predict population responses to environmental change?

Chevin (2015) and Janeiro et al. (in press) speculated that published IPMs that did not include explicit evolutionary processes could provide spurious insight. Three strands of evidence suggest this speculation may often be unwarranted.

First, the signature of evolutionary change in model predictions is a function of the heritability of the trait: when the phenotypic variance is dominated by the environmental component of the phenotype then the dynamics of that component will dominate model predictions. Most IPMs to date have been constructed for body weight (Merow et al., 2014), a trait that often has a heritability of less than 0.2 in vertebrates (e.g., blue tits; Garnett, 1981) and often around 0.1 (e.g., bighorn sheep; Wilson et al., 2005). This means that model predictions will be dominated by the dynamics of the environmental component of the phenotype and that a phenomenological statistical approach to parameterising these models has the potential to capture observed dynamics well.

Second, even when phenotypic traits are heritable, they rarely evolve in the wild as predicted: evolutionary stasis of heritable phenotypic traits in the presence of directional selection is frequently observed in nature (Merilä et al., 2001). When fitness functions are monotonic in the phenotypic value and selection is directional (which is typical for body size (Kingsolver et al., 2001)), then in order to maintain an equilibrium trait distribution the inheritance function must reverse the phenotypic changes caused by selection. Coulson and Tuljapurkar (2008) showed this for the mean phenotypic trait. However, when the genotype-phenotype map is additive and there is additive genetic variance for the trait, directional selection is expected to result in evolutionary change and the inheritance function for the genetic component of the phenotype can not reverse genetic changes attributable to selection. Unmeasured genetically correlated characters can prevent evolutionary change in these circumstances, although the cases when this is likely to prevent evolution are restrictive, and evidence for such characters playing a major role in limiting evolution in the wild is lacking (Agrawal and Stinchcombe, 2009). Assuming selection on the phenotype has been measured appropriately and is directional, this suggests that the assumption of an additive genotype-phenotype map may be violated, and the mean of the parental and offspring breeding value distributions may not be equal. A mechanism such as over-dominance can achieve this (Fisher, 1930). Our approach allows the effects of relaxing assumptions of quantitative genetics on evolutionary change to be approximated through the use of phenomenological inheritance functions for the genetic component of the phenotype.

Third, because evolutionary change is rarely observed in the wild when it is predicted, observed phenotype change in natural populations is usually attributable to plasticity (e.g. Ozgul et al., 2010, 2009). In these cases, standard, non-evolutionarily explicit, IPMs have accurately captured observed dynamics (Childs et al., 2003*a*; Merow et al., 2014; Ozgul et al., 2010).

These three strands of evidence suggest that evolutionarily explicit IPMs may frequently not be required to gain useful insight into population responses to environmental change. If there is no statistical evidence of temporal trends in inheritance, development or fitness function parameters once variation in the ecological environment has been corrected for, then the use of evolutionarily explicit IPMs may result in the construction of unnecessarily complex models. There is often a temptation to include ever more complexity into models, but this comes at the cost of analytical tractability: as more mechanisms or processes are incorporated into models, understanding why a model produces the predictions it does becomes increasingly challenging. However, when evolutionary change is convincingly documented (e.g. Reznick et al., 1997) or is proposed to be a possible mechanism generating rapid phenotypic change (Coltman et al., 2003), the construction of evolutionarily explicit IPMs is advised as the models allow separation of the roles of adaptive and plastic responses to environmental change.

We have shown how evolutionarily explicit IPMs can be constructed, invalidating the criticisms of Chevin (2015) and Janeiro et al. (in press) that IPMs have not been developed to incorporate the character-state approach of quantitative genetics. IPMs that are not evolutionarily explicit have been used to address many questions in ecology and their application has proven insightful (Merow et al., 2014). They are likely to remain widely used and we expect this use to result in important new insights. However, we have extended their utility to cases where evolutionary processes are known, or proposed, to be drivers of phenotypic change.

## Conclusions

In this paper we have developed a theoretical modeling approach that links demography and quantitative genetics to explore how populations will respond to environmental change. The approach is general, providing formal links between ecology and evolution. Our work builds upon a growing literature of developing evolutionarily explicit structured population models. This body of literature shows how flexible IPMs are. They provide a powerful tool with the potential to unify ecology and evolution.

## Acknowledgements

Bernt-Erik Saether, Luis-Miguel Chevin and Michael Morrissey participated in useful discussion and provided comments on an earlier version of the manuscript. Stephen Proulx, Thomas Reed, and two anonymous reviewers provided critical feedback that greatly improved the first submitted version of the paper. TC acknowledges support from NERC (grant number NE/K014218/1) and support of Centre for Advanced Study in Oslo, Norway, that funded and hosted the research project (Climate effects on harvested large mammal populations) during the Academic year of 2015/16.

## Appendix A: Model parameters and predicted dynamics

Model Parameterization

**Model A**:

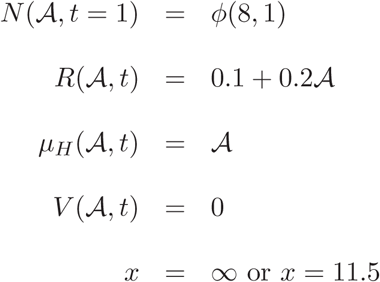

**Models B** and **C**:

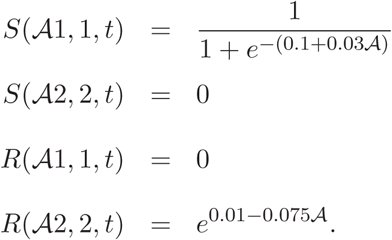

Starting conditions at time *t* = 1 are multivariate normal with the following parameters, Model B:

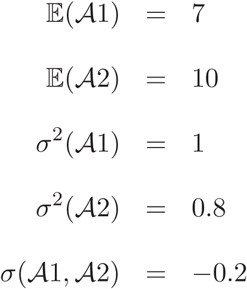

**Model C**:

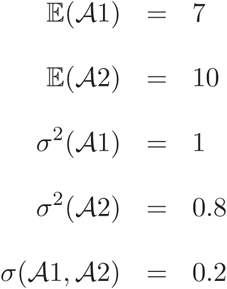

**Model D**:

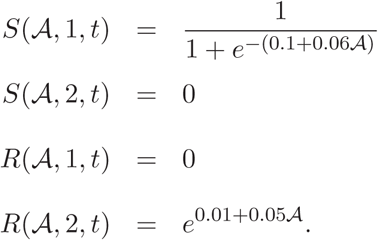

Starting conditions at time *t* = 1 for **model D:** 
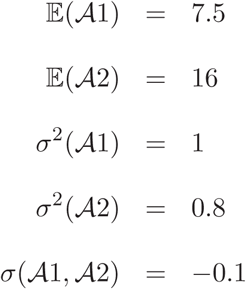

**Model E**

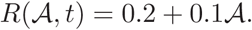

**Model F**: no plasticity:

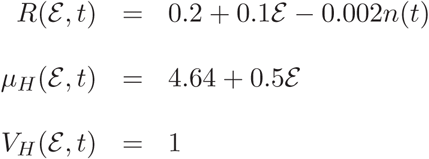

**Model G**: Adaptive phenotypic plasticity:

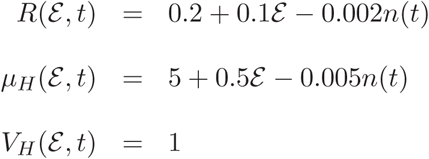

**Model H**: Non-adaptive plasticity:

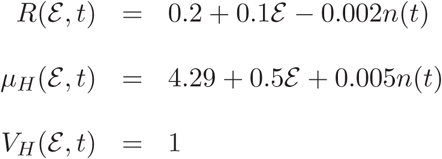

**Model I**

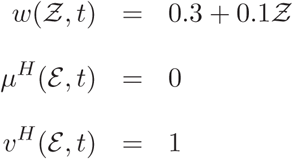

**Model J**

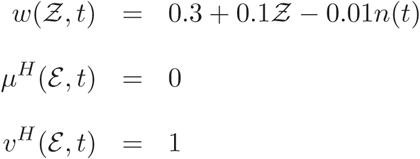

**Model K**

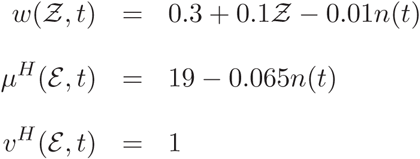

Initial starting conditions for 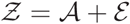 for **models I to K**:

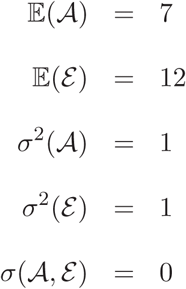

**Model L**

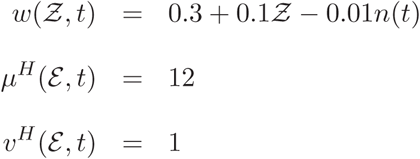

**Model M**

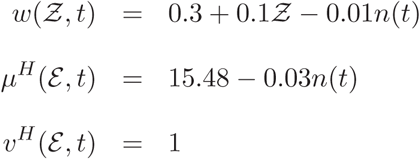

**Model N**

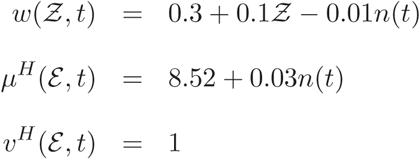

Initial starting conditions for 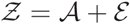 for **models L to N**: 
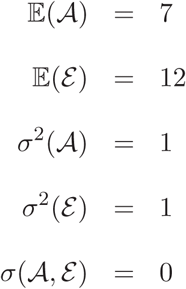
**Models P and Q**: 
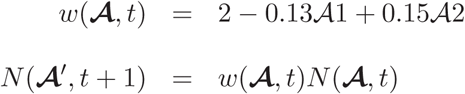
Starting conditions at time *t* + 1 for **model P**: 
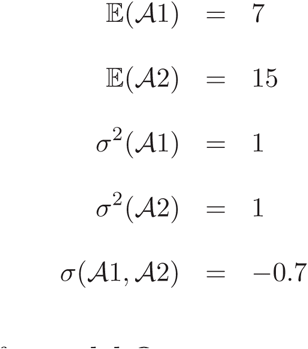
Starting conditions at time *t* + 1 for **model Q**: 
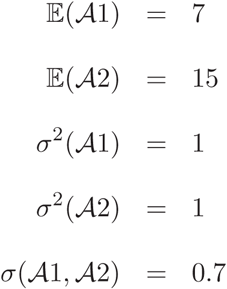

### Calculating quantities from model outputs

The expectation of a distribution of 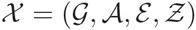 can be calculated as

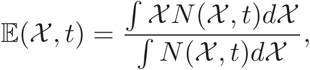

The variance of a distribution can be calculated as

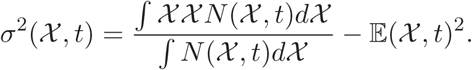

For a bivariate distribution 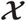 consisting of traits 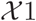 and 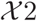 then the covariance between these two traits will be,

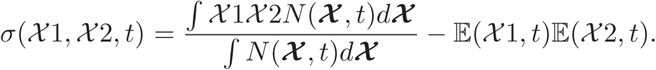

The skew can be calculated as,

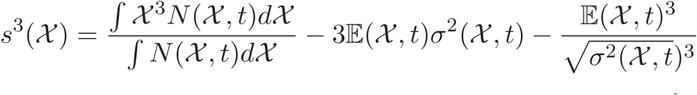

The kurtosis can be calculated in the following way. First, we define the n^th^ non-central moment of a distribution at time *t* as 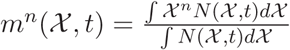 then,

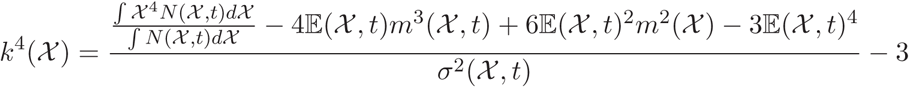

### Gaussian inheritance function when demography differs between males and females

The distribution of mothers and fathers at time t is respectively defined as 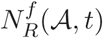 and 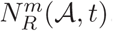. These distributions are the same size.

We can write 
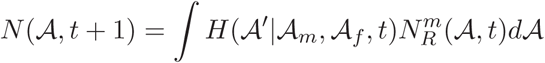
where the component functions of 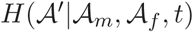 are 
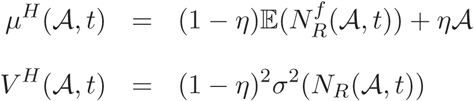
and 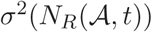 is the variance in 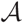 across all parents.

Alternatively, 
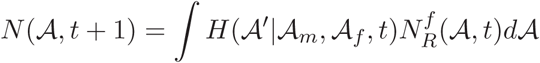
where the component functions of 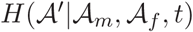 are 
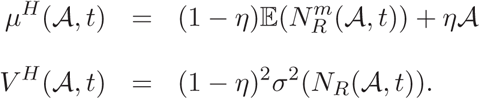

As the distributions 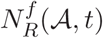 and 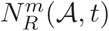 depart from normality, the approximations will predict dynamics that diverge from those predicted by the convolution.

### How do different functions alter character distributions?

Assume 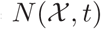 is proportional to a Gaussian distribution. The following parameterizations of a transition functions 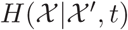 in a model 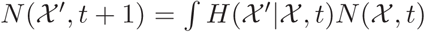 will have no effect on the location or shape of the distribution such that 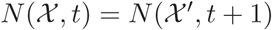

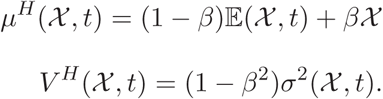

Note that in this model there is no fitness function and no selection.

When the intercept of 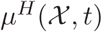 is less than 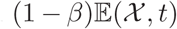 then 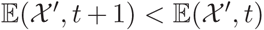 and vice versa. A function 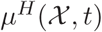 can consequently be parameterized to reduce the mean of a distribution across generations or time steps if desired.

The slope *β* will reduce 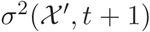 by *β*^2^ compared to 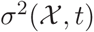. The intercept of 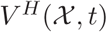 injects additional variation. If the intercept is larger than 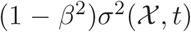 then 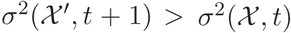. Functions 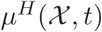 and 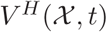 can consequently be selected to alter the variance from one time step or age to the next.

The further the distribution 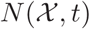 departs from normality, the more approximate these equalities will become. However, large departures from these equalities can be used to increase the mean or variance of any distribution in a desired direction.

In Figure A1 we show how 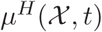 and 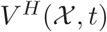 can be parameterized to modify the mean and variance of 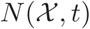 when it is proportional to a normal distribution.

**Figure A1.**
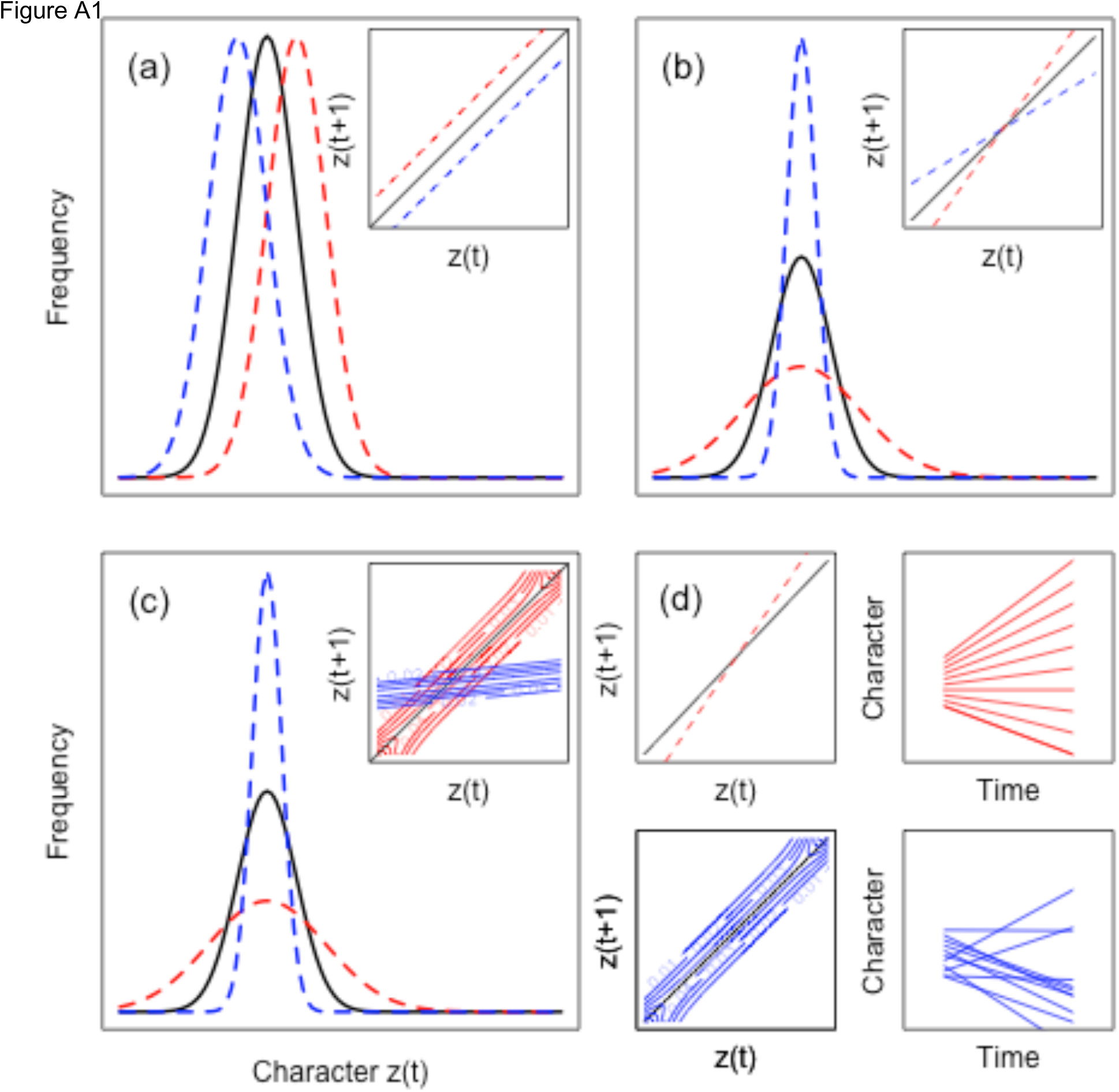
How parameterizations of transition functions for the environmental component of the phenotype 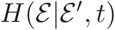 can be used to grow, maintain or shrink the mean and variance of 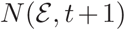. We start with a normal distribution. The initial distribution is represented with a black line in the main figures. The inset figures in (a) to (c) shows the transition functions, with the black line representing the function that has no effect on the location or shape of 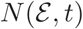. (a) increasing or decreasing the intercept of 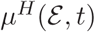 influences the location, but not the shape of 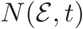. (b) How altering the slope of 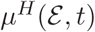 influences the shape of 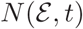. In this example the mean is unaffected as the function passes through the *x,y* co-ordinate 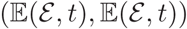. (c) how altering the intercept of 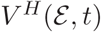 influences the variance. The transition functions in the insets of (b) and (c) generate distributions with the same means and variances (compare blue, red and black distributions in (b) and (c)). A change in variance between 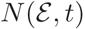 and 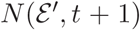 achieved by altering the slope of 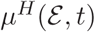 or the intercept of 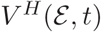 generates different amounts of mixing. In (d) upper and lower 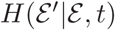 functions impact the variance to the same extend (left hand figures) except the red function simply spreads out the distribution without altering the relative rank of each individual. In contrast, the blue function changes relative ranks (right hand figures).

**Figure A2.**
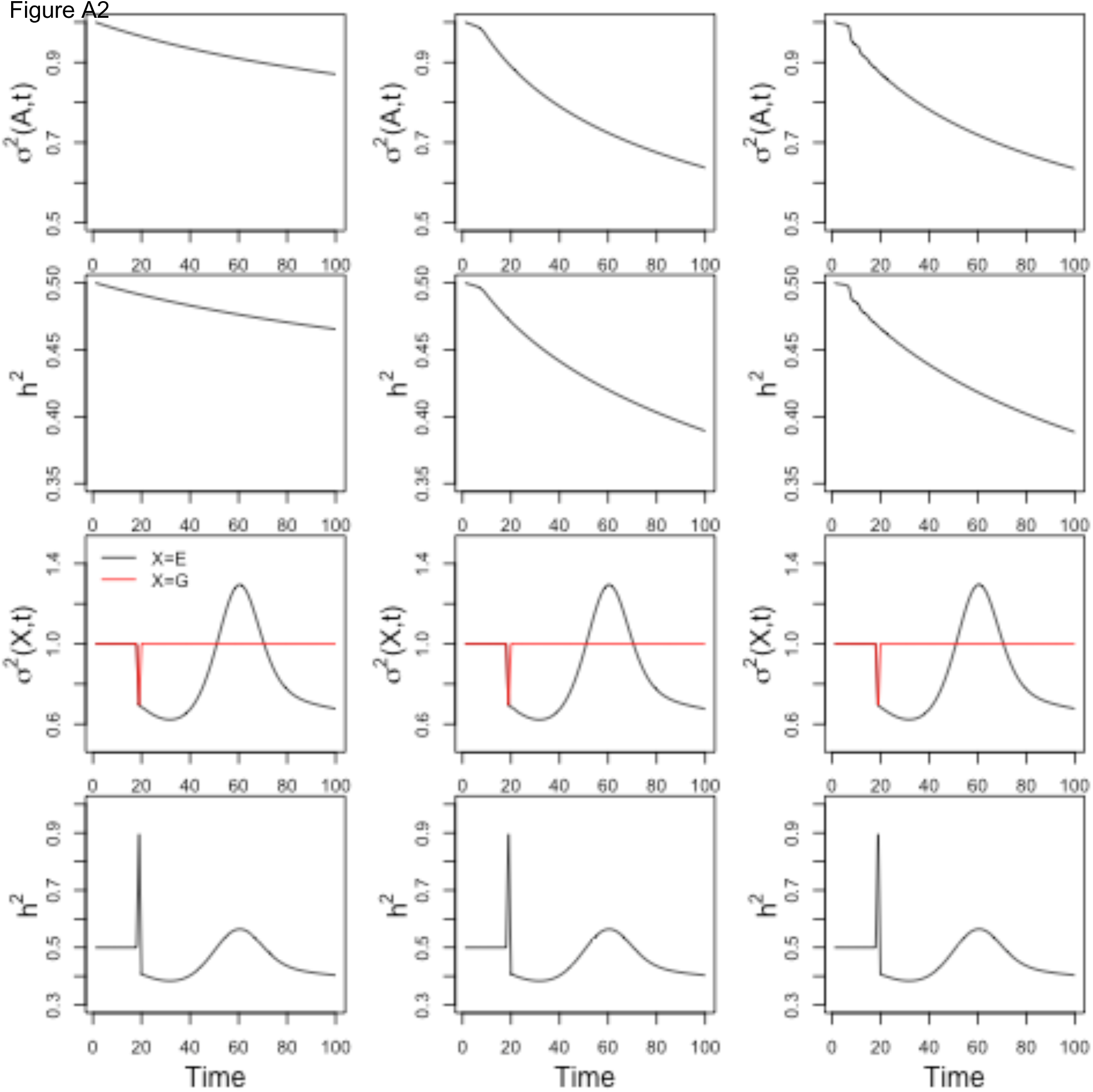
Dynamics of the additive genetic variance (a)-(c) and the heritability (d)-(f) in models I to K. Models of the additive genetic (back line) and environmental (red line) variance (g)-(i) and the heritability (j)-(l) in models L to N. See Figure 5 main paper for dynamics of means and population growth.

### Mortality selection and changes in the mean phenotype

When a trait is normally distribution, selection needs to be strong in order to substantially shift the mean of a phenotype distribution. Such strong selection inevitably leads to a decrease in population size. In Figure A3 we show how killing 25% of the heaviest individuals has only a small effect on the mean for a distribution with a mean of 0 and a standard deviation of unity. The evolutionary response is even less if 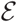 and 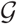 are uncorrelated. For example, in the example in Figure A3, the evolutionary response would be half the phenotypic response for *h*= 0.5. In order to substantially shift the mean of the a normal distribution via mortality selection it is necessary for the majority of the population to die.

**Figure A3.**
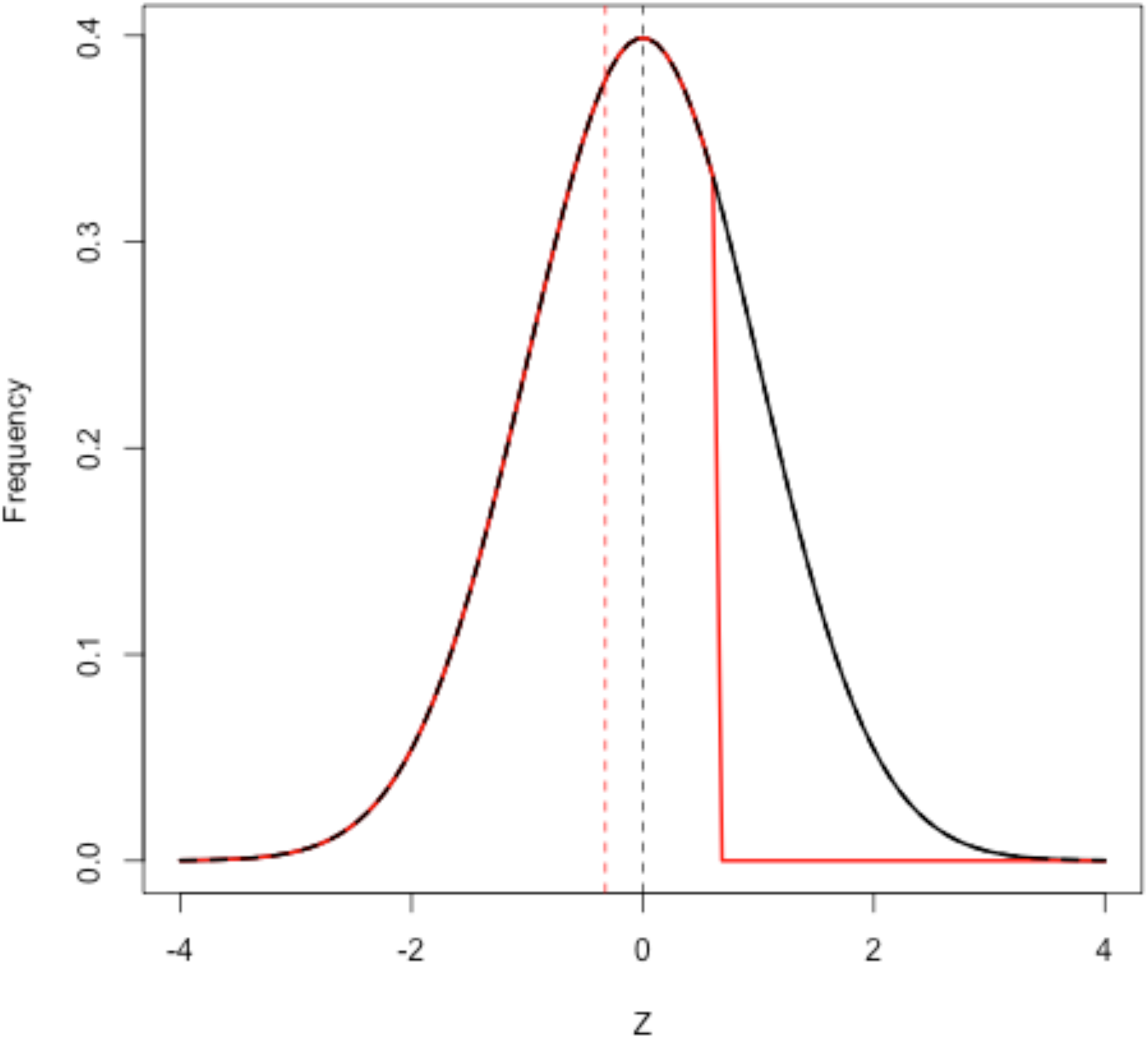
A normal distribution with mean 0 and standard deviation 1 prior to mortality selection (black line). Mortality occurs, killing off the top 25% of individuals (red distribution). The mean changes from 0 (vertical dashed line) to −0.0324. In other words, even a large highly selective mortality event has a relatively small effect on the mean of a normal distribution.

